# Circadian and Sleep-Wake Modulation of Functional Connectivity Across Brain Oscillations and States Linked to Cognition in Humans

**DOI:** 10.1101/2025.10.20.683464

**Authors:** Alpar S Lazar, Zsolt I Lazar, Nayantara Santhi, June C Lo, John A Groeger, Derk-Jan Dijk

**Affiliations:** Faculty of Medicine and Health Sciences, University of East Anglia, UK; Faculty of Physics, Babes-Bolyai University, Romania; Department of Psychology, University of Northumbria, UK; Centre for Sleep and Cognition, Human Potential Translational Research Programme, and Department of Medicine, Yong Loo Lin School of Medicine, National University of Singapore, Singapore; Department of Psychology, Nottingham Trent University, Nottingham, United Kingdom; Surrey Sleep Research Centre, Faculty of Health and Medical Sciences, University of Surrey, Guildford, UK; UK Dementia Research Institute, Care Research& Technology at Imperial College London and the University of Surrey Guildford

**Author notes:** **Corresponding author:** Alpar S Lazar **Email:**. **Author Contributions:** D.-J.D. and J.A.G. designed research; D.-J.D. directed the research; A.S.L., N.S., and J.C.L. performed research; A.S.L., and Z.I.L. analyzed data; and A.S.L., Z.I.L., N.S., J.A.G., J.C.L., and D.- J.D. wrote the paper. **Competing Interest Statement:** Derk-Jan Dijk is a consultant to Boehringer Ingelheim, Astronautx and Danisco Sweeteners, and collaborates and/or has received equipment from SomnoMed and VitalThings.

**Keywords:** EEG, Brain State, Homeostasis, Neural Networks, Phase Coupling

## Abstract

Sleep and circadian rhythms both contribute to cognitive performance, but the underlying neuronal network-level changes remain unclear. We quantified the contribution of brain state, sleep-pressure dynamics across the sleep-wake cycle, and circadian rhythmicity to electroencephalographic (EEG) functional connectivity (FC) and examined how these network changes relate to cognition. Thirty-four healthy adults completed a 10-day forced-desynchrony protocol to uncouple sleep-wake and endogenous circadian rhythms. From over 1,200 hours of artifact-free EEG, we derived phase-coupling metrics to quantify FC across brain states, thirds-of-the-night (sleep pressure), and circadian phase, and related these network measures to a range of cognitive performance indices. FC differed markedly between brain states, especially in the alpha and sigma bands, and was modulated by sleep history and circadian phase. Principal component analysis revealed both a global and a topographically distributed FC component which responded differentially to sleep pressure. Dissipation of sleep pressure was accompanied by increasing global FC in NREM sleep, and especially in the delta, sigma and beta frequencies, and decreasing global FC in the alpha band. During REM sleep, global FC decreased in nearly all frequency bands with dissipation of sleep pressure. The influence of circadian phase on FC was smaller than that of sleep pressure and varied across brain states. Lower global theta-band FC in NREM and alpha-band FC during wake predicted better alertness and working memory accuracy, an effect modulated by circadian phase. These results suggest that sleep homeostasis and circadian timing interact to stabilize functional brain connectivity in wakefulness, thereby supporting optimal cognitive function.

**Significance Statement:** Identifying how sleep restores neural networks degraded during wakefulness is of significance for understanding sleep’s role in maintaining brain function. Here we used a protocol to isolate effects of sleep from effects of circadian rhythmicity on functional connectivity of neural networks and assessed associations with cognitive performance. We found that circadian rhythmicity but in particular the dissipation of sleep pressure had profound effects on global connectivity which were different for NREM sleep, REM sleep and wakefulness and varied across frequency bands. Connectivity measures in the theta, alpha and sleep spindle frequency ranges were associated with cognitive performance. These novel findings provide a new perspective on the nature of the sleep recovery process contributing to the waking performance capability of human brains.

## Introduction

Sleep is widely conceptualized as a recovery process, which returns neural network function degraded by prior wakefulness toward a baseline state (1). Recovery may be understood mechanistically (e.g. synaptic downscaling) or functionally (e.g. restoration of information transfer and integration capability); yet exactly how recovery unfolds across sleep and circadian cycles remains poorly defined.

Oscillations in the electrical activity of the cortex are thought to reflect the fundamental neurophysiological mechanisms enabling the temporal organization, transfer, and integration of information in the brain which are essential for waking performance (2–5). Tracking how these rhythms are restored or reshaped during sleep may reveal the network mechanisms of recovery.

In humans, this oscillatory activity can be assessed by intracranial recordings of unit activity or field potentials and noninvasive techniques such as magnetoencephalography and electroencephalography (EEG) (6, 7). The power spectrum of the EEG signal derived from single electrode sensors provides information on the frequency composition of the oscillatory activity at the location of the sensors and the underlying cortical neural firing rate and synchronicity (2). The frequency composition and amplitude of the EEG vary across and within brain states (wakefulness, NREM sleep, and REM sleep)(8). Furthermore, both the sleep-wake cycle and circadian rhythmicity have been shown to modulate brain oscillatory activity as quantified by power spectral analysis within each of the brain states (9, 10). Among the most prominent modulations within vigilance states are the decline of slow wave activity within NREM sleep from the beginning to the end of sleep episodes and an increase of theta activity during wakefulness with elapsed time awake (11, 12). These dynamics of low-frequency EEG activity, which are observed in many mammalian species, have been used extensively as indicators of a recovery process during sleep and a buildup of sleep pressure/sleep debt and associated deterioration of brain function during prolonged wakefulness (13). However, it has been challenging to establish how the dynamics of slow wave activity relates to a sleep dependent recovery process underpinning brain function during wakefulness.

Beyond the frequency distribution of the EEG signal measured by the power spectrum, the functional integration of brain regions can also be characterised using connectivity metrics. A widely used measure is the phase-lag index (PLI) (14) which measures the consistency with which the phase of a signal from one location leads or lags a signal from another location by ignoring phase differences centred around zero. By taking the sign of the phase difference and averaging over time, PLI isolates stable, non-instantaneous phase relationships that reflect genuine synchrony. A commonly used extension of the PLI is the weighted phase-lag index (wPLI), which not only discards phase differences around zero but also weights each observation by the magnitude of the imaginary component of the cross-spectrum (Vinck et al., 2011). By emphasizing larger phase-lag magnitudes, wPLI further suppresses the influence of small, noise-driven phase fluctuations and residual volume-conduction effects that can still bias the unweighted PLI. In practice, wPLI therefore provides a more robust indicator of true, non-instantaneous synchrony, and hence potentially causal interactions, between cortical areas, strengthening our inferences about information transfer and integration across distributed networks. A further refinement of this connectivity metric is the debiased weighted phase lag index (dwPLI) which corrects for sample-size bias in wPLI and further reduces noise-driven inflation of connectivity estimates. This yields a more stable and interpretable measure of non-instantaneous functional connectivity (see Methods section).

EEG connectivity measures are often used in cognitive neuroscience research to investigate associations between functional connectivity (FC) and brain function at single time points. Several studies have identified changes in FC in specific networks, such as the attentional or default mode networks, and their associations with performance on tasks that depend on these networks. Changes in FC are also associated with brain development and brain disorders, such as mild cognitive impairment (MCI) and Alzheimer’s disease (15–19).

EEG connectivity measures also vary across brain states, indicating that they do not just reflect anatomical/structural connectivity but FC (15, 20, 21). One argument for the functional importance of connectivity measures is the observation that the reduced responsiveness to external stimuli associated with sleep is accompanied by decreased functional brain connectivity (22).

Whether FC measures of brain oscillatory activity in the various brain states are modulated by dissipation and increase of sleep pressure and circadian rhythmicity has not been established. Finally, it has not been established whether and how functional EEG connectivity measures relate to the well-established sleep-wake and circadian rhythmicity-dependent modulation of brain function as quantified by performance on a variety of cognitive tasks (23–25).

We aimed to clarify these questions and hypothesized that the physiological effects of both sleep pressure (its dissipation and accumulation) and circadian phase would influence functional connectivity (FC). We also hypothesize that FC will be linked to waking cognition. Given that brain oscillatory activity exhibits brain-state (Wake, NREM, and REM) and frequency-specific patterns, we further hypothesized that these effects would be specific to particular brain states and frequency bands.

The separate contribution of the sleep-wake cycle and circadian rhythmicity-driven process to the restoration and deterioration of brain function cannot be assessed by analysis of sleep and wakefulness under baseline conditions because under these conditions these processes are entangled. Quantifying their separate contribution requires desynchronizing the sleep-wake cycle from endogenous circadian rhythmicity driven by the suprachiasmatic nuclei and indexed by melatonin, cortisol, and core body temperature rhythms (10, 11, 23, 26).

Here, we quantified the independent contribution of circadian rhythmicity, and the dissipation and accumulation of sleep pressure (i.e., time elapsed in sleep and wake, respectively) to functional brain connectivity (FC) as measured by an improved index of phase synchronization, i.e., debiased weighted phase leg index (dwPLI) (27) in both sleep and wake and its association with cognition.

FC was quantified across more than 1,200 hours of artifact-free NREM and REM sleep and nearly 90 hours of wake EEG, alongside performance on tests of vigilance and working memory, and processing speed. The data were collected in a 10-day forced-desynchrony protocol in 34 healthy young adults (Fig. 1).

**Fig 1.**
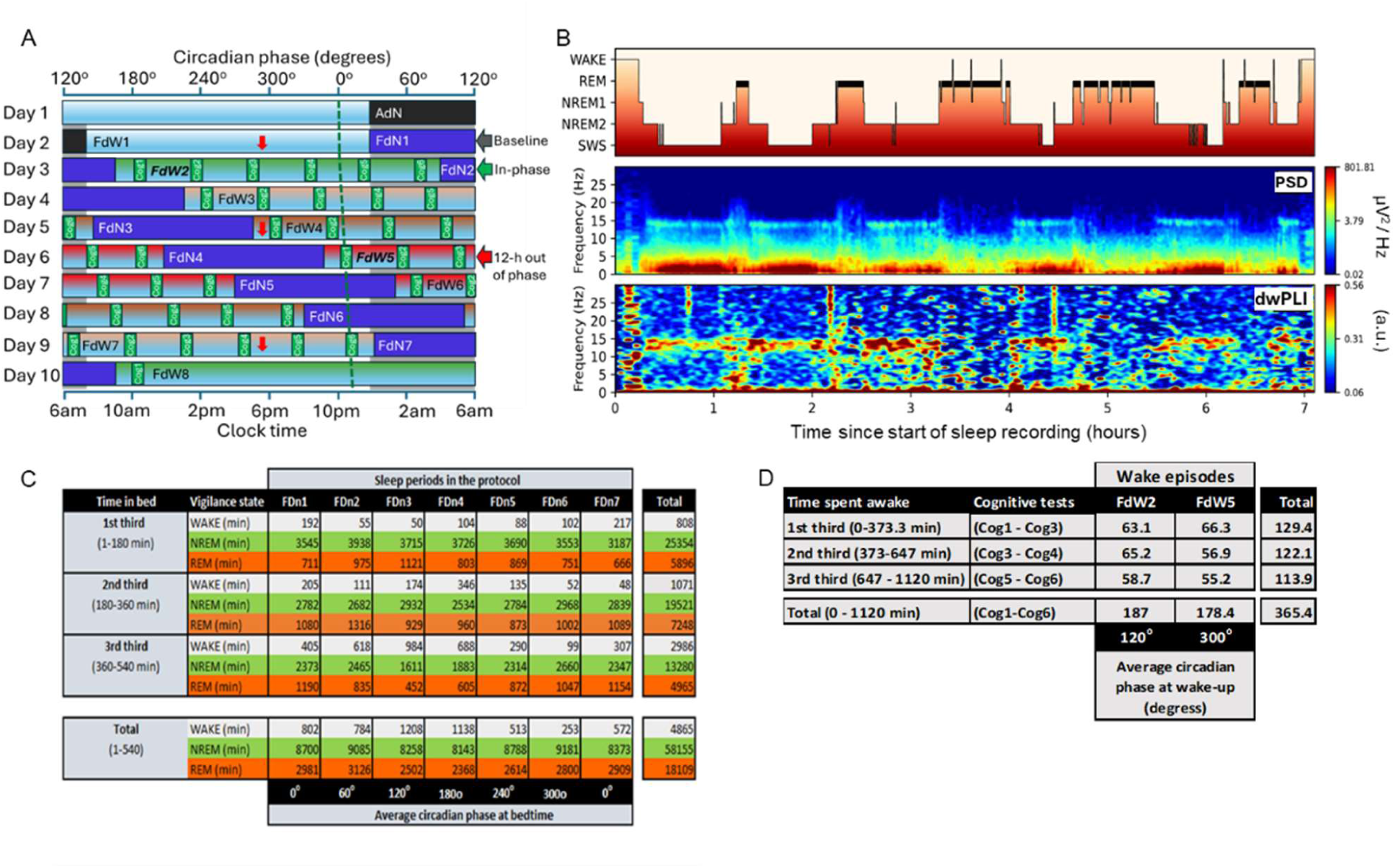
Research protocol and EEG data analysis. (A) Raster plot of the 28-h forced desynchrony (FD) protocol with a representative example of sleep-wake schedule (habitual bedtime: 00:00). Following an 8-hour adaptation night (ADn) and adaption day (FdW1) participants were scheduled to a 28-h sleep-wake cycle, including 9 hours and 20 minutes of sleep (horizontal dark blue bars) and 18 hours and 40 minutes for wakefulness (horizontal light blue bars). The 24- h variation of plasma melatonin concentration was assessed on three occasions, at baseline (FdW1-FDn1), FdW4- FdN4, and FdW7-FdN7 to estimate the phase and period of the intrinsic circadian clock as indicated by the GREEN line, which represent the estimated onset of melatonin. The red arrow pointing indicates the start of each 24-hour-long melatonin sampling session. Vertical green bars indicate cognitive test sessions included in the current analysis. The gradient color on the wake period, ranging from green (habitual timing) to red (12 h out of phase), indicates circadian alignment of the wake periods with the FD protocol. The green and red arrows to the right of the figure indicate the two wake periods (FdW2 and FdW5) included in the wake EEG analysis. During these two wake periods, each cognitive test session included two 2-minute Karolinska Drowsiness Tests (KDTs) to measure resting wake EEG at the start and end of the assessments. (B) Time course of sleep stages (top panel), power spectral density (PSD) (middle panel), and debiased weighted phase lag index (dwPLI) (bottom panel) measured during a baseline sleep episode in one participant. Warmer colors represent higher spectral power (logarithmic scale) and functional connectivity (FC) as measured by dwPLI. Temporal resolution is 10 seconds/pixel, each representing the average of the short-time Fourier transforms of four segments of 4-s long, Hanning tapered windows with 50% overlap. Power spectral density (PSD) values were obtained by also averaging over the 12 channels (FP1, Fp2, F3, F4, C3, C4, T3, T4, P3, P4, O1, and O2) whereas dwPLI values were averaged over all 66 electrode pairs. (C) The amount of artifact-free Wake, NREM, and REM sleep EEG data (in minutes) analyzed for each sleep episode throughout the 10-day-long forced desynchrony study. (D) The amount of artifact-free Wake EEG data (in minutes) analyzed for the baseline (FdW2) and the 12-h out-of-phase (FdW5) wake episodes.

We conducted a comprehensive analysis involving principal component analysis (PCA) applied to the functional-connectivity (FC) matrices derived from 66 electrode pairs and 21 cognitive outcome measures. We found a global principal component (PC), reflecting widespread synchronization (global FC). The second PC reflected a topographically distributed connectivity (distributed FC). To relate these network patterns to cognitive function, we conducted a PCA on the cognitive measures and analyzed the first two cognitive PCs representing cognitive alertness and working memory performance (PC1) as well as processing speed and working memory performance (PC2).

We found that recovery of brain function during sleep is associated with an increase in global FC during NREM sleep and a decrease during REM and to some extent in wakefulness during spontaneous interruptions of sleep. The circadian day was associated with a reduction in global FC during wakefulness in the alpha band, and this reduction associated with better cognitive performance. Together, these data illuminate how sleep homeostasis and circadian timing orchestrate the recovery of large-scale network connectivity, enabling optimal brain function upon waking.

## Results

### Effect of brain states, homeostatic sleep pressure, and circadian phase on functional EEG connectivity spectra

We used linear mixed effects modelling to quantify the effects of the three consolidated brain states – NREM, REM sleep, and wake – during the sleep episodes as well as the effects of time elapsed in sleep (i.e., dissipation of sleep pressure) and circadian phase on power spectrum density (PSD) and EEG connectivity measures (Fig. 2). NREM included NREM2 and slow wave sleep (SWS) following common practice (Dijk at al. 1995, Lazar et al., 2015). Artifact-free EEGs recorded during 231 in bed periods of 9h and 20 mins each, which were distributed across the circadian cycle, were analyzed for 12 electrodes (Fp1,Fp2,F3,F4,C3,C4, P3,P4,T3,T4,O1,O2) and 66 electrode pairs (combinations of 12 electrodes) and frequency bins between 0.5 and 32 Hz (Figs 1*A*-*C*). EEG connectivity was estimated using the debiased weighted phase lag index (dwPLI), as well as additional PC metrics including weighted phase lag index, phase lag index, imaginary coherence, and phase coherence (Fig S1). In a first analysis step we averaged the absolute power spectrum density (PSD) across the 12 EEG channels and connectivity metrics across all 66 individual intra and interhemispheric electrode pairs.

**Fig 2.**
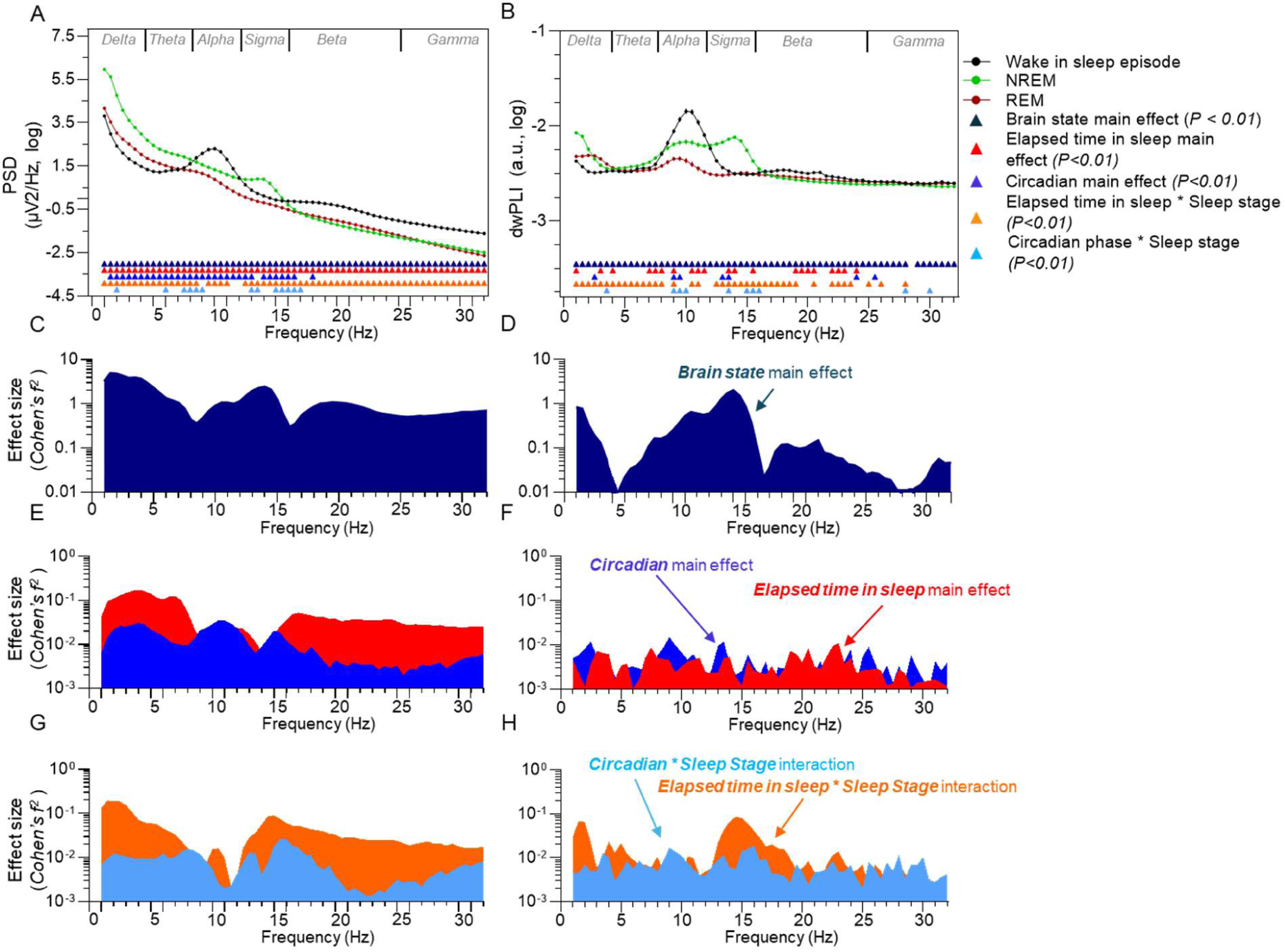
Power spectral density and functional connectivity spectra across brain states and sleep episodes. The main effect of brain states on (A) power spectrum density (PSD) and (B) debiased weighted Phase Lag Index (dwPLI) across all sleep episodes (231 nights) and averaged across EEG channels and cannels pairs, respectively. The upper panels represent least-square means (Lsmeans) and standard error of the mean (SEM). (C and D) Effect size (Cohen’s f^2^) of main effects of brain state on the PSD and dwPLI. (E and F) Effect size of main effects of Elapsed time in sleep (red) and Circadian phase (blue) on PSD and dwPLI. (G and H) Effect size of interactions between factors Elapsed time in sleep and Circadian phase on PSD and dwPLI. Colored triangles in (A) and (B) indicate significant (P<.01) effects. Detailed statistical results are in Dataset S1.xlsx.

PSD averaged across EEG channels showed the known characteristics of brain states (Fig 2*A* and Dataset S1.csv). Alpha (8-12 Hz) activity was higher in wakefulness than in NREM and REM sleep; delta (0.5-4 Hz), theta (4 - 8 Hz), and sigma (12-16 Hz) activity in NREM exceeded the corresponding values in REM.

The dwPLI also varied significantly across brain states and frequency (Fig. 2*B*, Dataset S1.xlsx). Its spectral profile was characterized by peaks in the alpha and sigma bands. Unlike PSD, dwPLI values did not markedly decrease with increasing frequency. Brain state effects on dwPLI were most dominant in the alpha-sigma range as well as in the delta frequencies (Fig 2*C*). Within the alpha band, dwPLI values were highest during wakefulness, lowest in REM sleep, and intermediate in NREM. In the sigma range (12– 16 Hz), dwPLI peaked most strongly during NREM 2 and SWS. Overall, brain states were best distinguished by PSD in the delta and sigma bands (Fig. 2*A,C*) and by dwPLI values in the alpha and sigma bands (Fig. 2*B,D*). In a confirmatory analysis, we repeated this procedure using four additional FC metrics (imaginary coherence, phase coherence, phase-lag index, and standard weighted phase-lag index and observed nearly identical brain state-dependent spectral profiles, confirming that our findings are robust to the choice of FC measure (Fig S2*A-E* and Dataset S1.xlsx). For exploratory purposes, we also repeated the same analysis including all NREM substages in the model (NREM1, NREM2, SWS as well as REM and wake during sleep) (Fig S3 and Dataset S2.xlsx).

Mixed model analyses revealed significant (p<0.01) main effects of Time Elapsed in Sleep and Circadian Phase on PSD across all frequencies (Fig. 2*E*, Dataset S1.xlsx). For dwPLI these effects were much smaller and restricted to specific frequency bands (Fig. 2*F*, Dataset S1.xlsx). The sleep-pressure and circadian effects on dwPLI were substantially smaller than the effects of Brain State (Fig. 2*D* and *F*). For PSD, the strongest sleep-pressure effects occurred in the delta and theta bands, reflecting the well-known decline of low-frequency power during sleep, while the largest circadian modulations peaked in the alpha (8-12 Hz) and sigma bands (Fig. 2*E*). Except near 10 Hz and 15 Hz, sleep-pressure effects on PSD exceeded circadian effects. For dwPLI, sleep-pressure effects on connectivity were largest in the theta, and beta bands, with magnitudes similar to or larger than circadian effects (Fig. 2*F* and Dataset S1.xlsx). Circadian effects on dwPLI peaked in the low delta (0.5–3 Hz) and a narrow sigma band (∼13 Hz). Overall, dwPLI was less strongly influenced by circadian phase than elapsed time in sleep. The interaction between brain state and both time elapsed in sleep and circadian phase showed more similar effect magnitudes across PSD and dwPLI (Fig. 2*G* and *H*). This suggests that while sleep pressure and circadian phase may not independently affect FC, their influence is strongly modulated by brain state.

A confirmatory analysis using four additional connectivity metrics, imaginary coherence, phase coherence, wPLI, and PLI, produced effect-size profiles for elapsed time in sleep and circadian phase, comparable to those observed with dwPLI (Fig S2*F and* Dataset S1.xlsx). Some metrics, particularly imaginary coherence and phase coherence showed steeper sleep-dependent effect sizes and smaller circadian effects (Fig S2).

Because dwPLI provided a good balance between sensitivity to state-related changes and resistance to volume-conduction artifacts, we retained dwPLI and excluded the other connectivity measures from further analyses (see methods). We also did not pursue further analysis of power spectral density (PSD), as these effects have already been well-documented in the existing literature (9).

### Principal components in the topographical distribution of functional connectivity

In the next step, we investigated the direction and topography of significant main effects of elapsed time in sleep and circadian phase on functional connectivity (FC). Given the large number of dependent variables (66 electrode pairs per brain state and frequency band), we first applied dimensionality reduction using principal component analysis (PCA) on FC as estimated by dwPLI from the 231 sleep episodes scheduled across the circadian cycle.

We conducted separate PCAs for each brain state NREM (Fig. 3), REM (Fig. 5), and wake during time in bed (Fig. 7) and each frequency band, resulting in a total of 18 analyses. Across all six frequency bands and three brain states, the PCA consistently identified one dominant component (PC1), which explained between 30% and 64% of the variance (mean: 41%) (Table S1.xlsx). The second principal component (PC2) accounted for 4% to 19% of the variance (mean: 9%). Overall, the combined variance explained by the first two components was highest in NREM (mean: 55%), followed by REM (mean: 49%) and wake during the sleep episode (mean: 45%) (Table S1.xlsx).

**Fig. 3.**
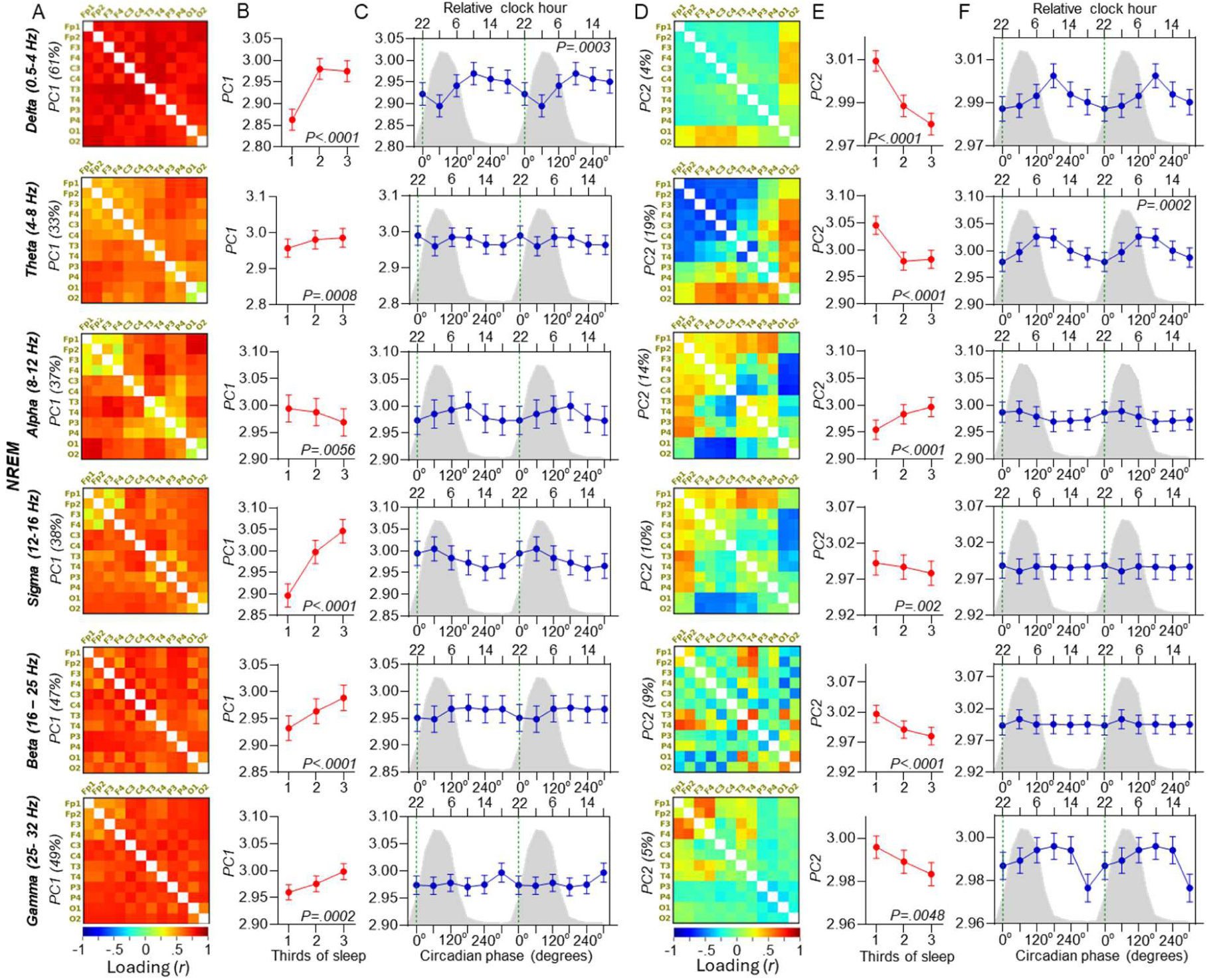
Principal component analysis of functional connectivity (dwPLI) during NREM sleep: effects of sleep progression and circadian phase. (A and D) Loading values of individual electrode pairs on PC1 and PC2, for delta, theta, alpha, sigma, beta and gamma frequencies. Values between brackets along the vertical axes represent the variance explained by the PC. (B and E) The modulation of PC1 and PC2 by elapsed time in sleep. Least square means (LSMeans) and standard errors of the mean (SEM) are presented for each 186.7-minute interval (i.e., one-third of the sleep episode), averaged across all studied circadian phases. (C and F) the circadian modulation of PC1 and PC2, with LSMeans and SEM shown for each 60° (∼4-hour) circadian phase bin, averaged across all sleep intervals. Data are double plotted to enhance the visualization of circadian rhythms. The green vertical line marks the dim-light melatonin onset (DLMO; 0° circadian phase), and the grey shaded area represents the average melatonin profile. Type III fixed effects are reported, with only statistically significant p-values (α = 0.01) indicated. Detailed statistical results are provided in Dataset S4.xlsx.

The correlation patterns between the principal components (PCs) and dwPLI values across individual electrode pairs indicated that higher loadings on PC1 reflected increased ‘global connectivity,’ characterized by overall elevated dwPLI across all electrode pairs (Figs. 3*A*, 5*A*, 7*A* and Dataset S3.xlsx). This pattern was consistent for PC1 across all frequency bands and brain states. In contrast, PC2 exhibited a more differentiated topography, and the interpretation of high loadings varied by frequency band and brain state (Figs. 3*D*, 5*D*, 7*D* and Dataset S3.xlsx). Accordingly, PC2 was interpreted as reflecting ‘topographically distributed connectivity.’ We limited our subsequent analyses to the first two principal components to optimize explained variance while controlling for Type I error. Although the second component explained a smaller proportion of variance, it was retained to mitigate the risk of Type II error.

### Effects of sleep-pressure dissipation on global and topographically distributed functional

We conducted two mixed model analyses: a primary analysis focusing on the two principal components (PCs) and an exploratory analysis examining connectivity in each electrode pair separately.

The number, magnitude, and direction of the significant (*P*<0.01) effects of elapsed time in sleep and circadian phase varied with brain state, frequency, and PC / topography (Figs. 3, 5, 7 and Dataset S4.xlsx).

As the sleep episode progressed and sleep pressure decreased, global FC (PC1) during NREM sleep markedly increased in most frequency bands, except for alpha, where it slightly but significantly decreased (Fig. 3*B*). In contrast, topographically distributed connectivity (PC2) decreased across all frequency bands except alpha, where it increased (Fig. 3*E*). The exploratory analysis supported these findings, revealing that most electrode pairs showed a significant (*P*<0.01) increase in coupling over the course of the sleep episode (Fig 4*A*, B and Dataset S5.xlsx). The widespread statistical significance of the effect across the studied EEG channel pairs is reflected in the small median p-value (median p = 1.91 × 10⁻¹¹) calculated among all comparisons with p < 0.01. In accordance with the results for PC1 (Fig. 3*B*) the electrode pair- based analyses showed that the delta and in particular the sigma but also the beta band exhibited the strongest and most widespread increases in FC with decreasing sleep pressure (Fig. 4*A*). Theta, alpha, and beta bands displayed a topographically split pattern, with some electrode pairs showing increases and others decreases in coupling (Fig. 4*A*). Notably, in the theta band, long-range fronto-centro-temporal connections exhibited an elapsed time in sleep-dependent increase in coupling, whereas long-range fronto-occipital and centro-occipital connections showed a decrease over the course of the sleep episode. In the alpha band, elapsed-time-in-sleep-dependent increases were predominantly observed in interhemispheric connections between homologous regions. Decreases in connectivity over the course of the sleep episode in the alpha band were primarily found in fronto-occipital and long-range fronto-centro-parietal connections (Fig. 4A and Dataset S4.xlsx).

**Fig 4.**
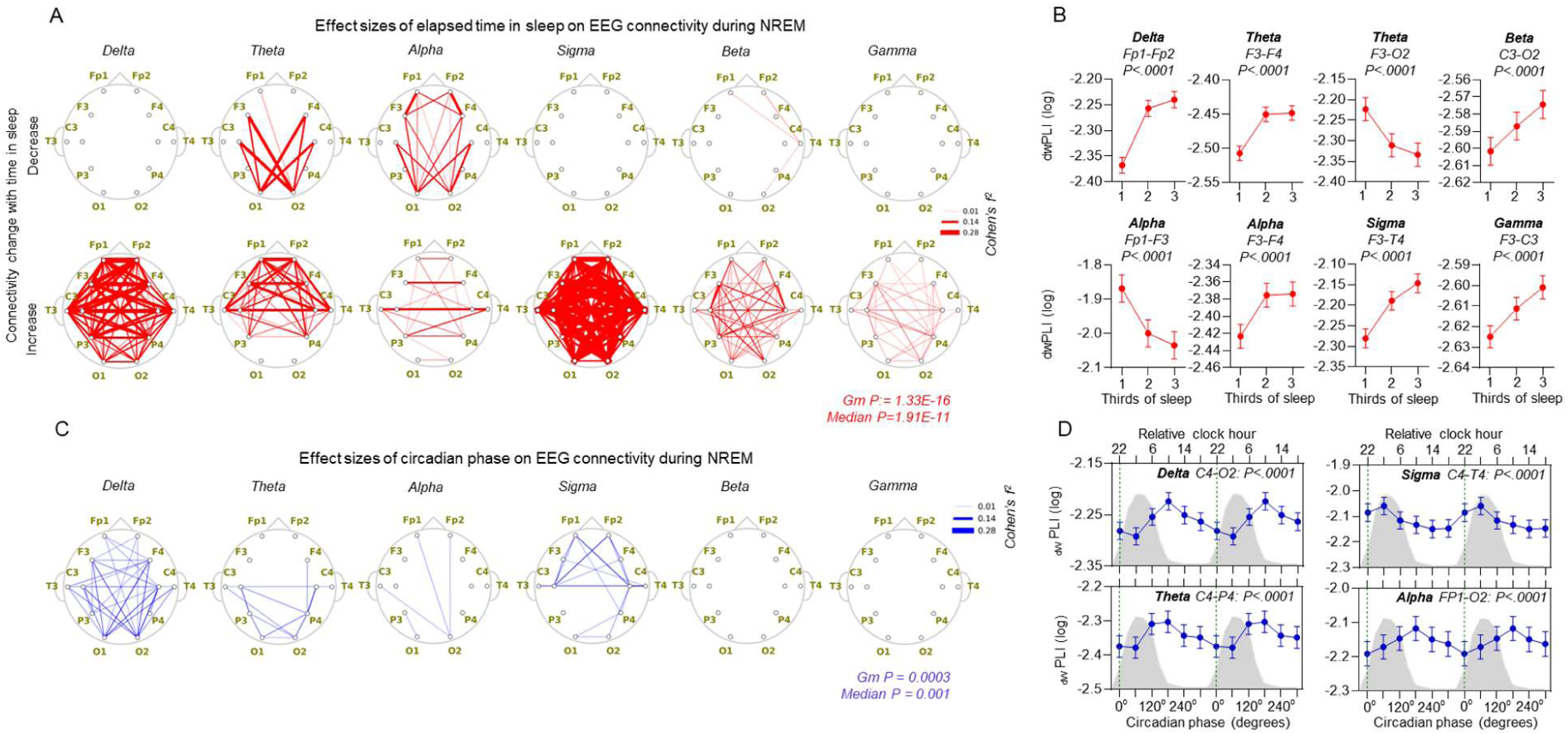
Topographical patterns of functional connectivity (dwPLI) in NREM sleep as a function of sleep progression and circadian phase. (A) Red lines represent the magnitude of effect sizes for elapsed time in sleep for electrode pairs, where a significant main effect of this predictor was observed (p < 0.01). Effect sizes for electrode pairs that showed a time-in-sleep–dependent increase or decrease in functional connectivity are shown separately. The geometric mean and median p-value among all significant p-values are also indicated and highlighted in red. (B) Representative examples of the effects of elapsed time in sleep on dwPLI. Least square mean (LSmeans) and standard error of the mean (SEM) are presented indicating sleep-dependent estimates at each 186.7-minute interval (third of the sleep episode) measured across all studied circadian phases. (C) Blue lines represent the magnitude of effect sizes for elapsed time in sleep for electrode pairs, where a significant main effect of this predictor was observed (p < 0.01). The geometric mean and median p-value among all significant p-values are also indicated and highlighted in blue. (D) Representative examples of the effects of circadian phase on dwPLI. LSmeans and SEM indicate circadian phase- dependent estimates at 60 degrees (∼ 4 h) bins measured across all studied sleep intervals and EEG derivations presenting a significant circadian modulation as shown in A. Data are double plotted to enhance the visualization of circadian rhythms. The green vertical line marks the dim-light melatonin onset (DLMO; 0° circadian phase), and the grey shaded area represents the average melatonin profile. Type III fixed effects are reported, with only statistically significant p-values (α = 0.01) indicated. Detailed statistical results are provided in Dataset S5.xlsx.

REM sleep (Fig. 5*B* and Dataset S4.xlsx), unlike NREM, showed a significant decrease in global FC (PC1) with elapsed time in sleep across most frequency bands, except for alpha, where no significant change was observed. In turn, topographical distributed FC (PC2) showed no significant effect of sleep pressure, except in the delta and beta frequency bands, where it increased with time elapsed in sleep (Fig. 5*E*).

The exploratory topographical analysis supported the PC1 findings showing that all electrode pairs that exhibited a significant elapsed-time-in-sleep-dependent effect (median *P =*5.91E-08) were characterized by a decreasing coupling over the course of the sleep episode (Fig. 6*A, B* and Dataset S5.xlsx). The largest number of electrode pairs showing a significant decrease in connectivity was observed in the sigma band.

**Fig 5.**
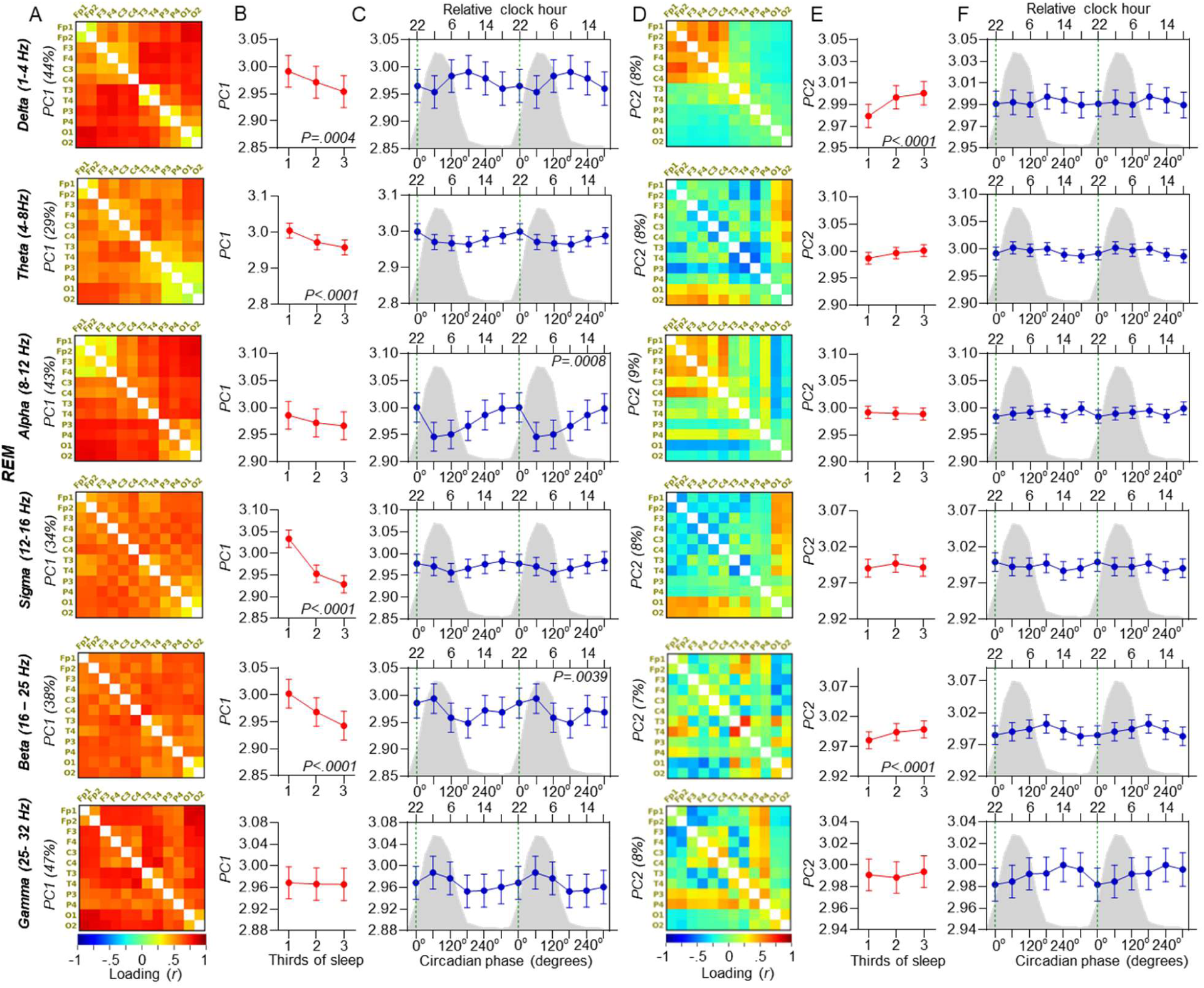
Principal component analysis of functional connectivity (dwPLI) during REM sleep: effects of sleep progression and circadian phase. (A and D) Loading values of individual electrode pairs on PC1 and PC2, for delta, theta, alpha, sigma, beta and gamma frequencies. Values between brackets along the vertical axes represent the variance explained by the PC. (B and E) The modulation of PC1 and PC2 by elapsed time in sleep. Least square means (LSMeans) and standard errors of the mean (SEM) are presented for each 186.7-minute interval (i.e., one-third of the sleep episode), averaged across all studied circadian phases. (C and F) the circadian modulation of PC1 and PC2, with LSMeans and SEM shown for each 60° (∼4-hour) circadian phase bin, averaged across all sleep intervals. Data are double plotted to enhance the visualization of circadian rhythms. The green vertical line marks the dim-light melatonin onset (DLMO; 0° circadian phase), and the grey shaded area represents the average melatonin profile. Type III fixed effects are reported, with only statistically significant p-values (α = 0.01) indicated. Detailed statistical results are provided in Dataset S4.xlsx.

**Fig 6.**
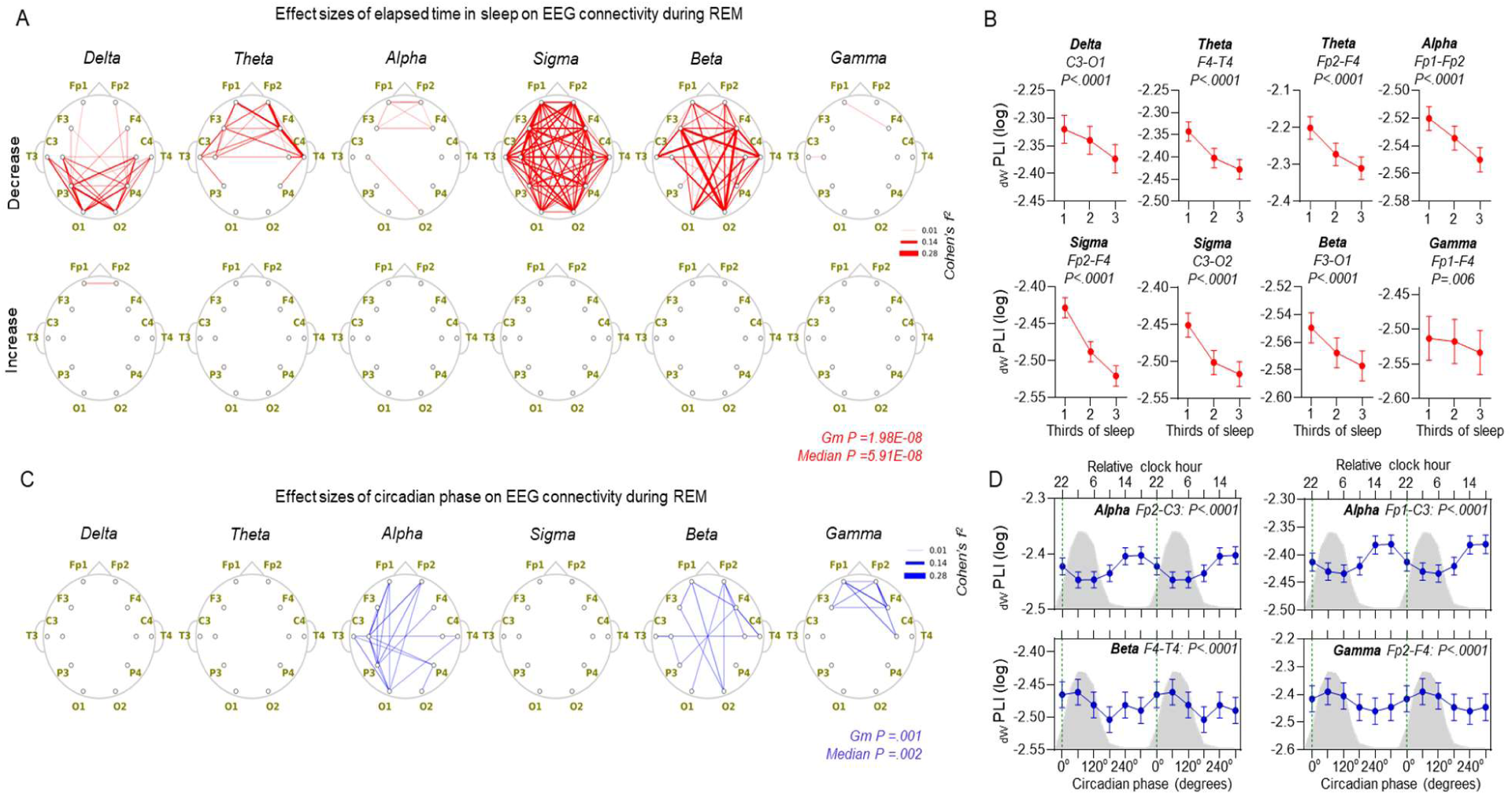
Topographical patterns of functional connectivity (dwPLI) in REM sleep as a function of sleep progression and circadian phase. (A) Red lines represent the magnitude of effect sizes for elapsed time in sleep for electrode pairs, where a significant main effect of this predictor was observed (p < 0.01). Effect sizes for electrode pairs that showed a time-in-sleep–dependent increase or decrease in functional connectivity are shown separately. The geometric mean and median p-value among all significant p-values are also indicated and highlighted in red. (B) Representative examples of the effects of elapsed time in sleep on dwPLI. Least square mean (LSmeans) and standard error of the mean (SEM) are presented indicating sleep-dependent estimates at each 186.7-minute interval (third of the sleep episode) measured across all studied circadian phases. (C) Blue lines represent the magnitude of effect sizes for elapsed time in sleep for electrode pairs, where a significant main effect of this predictor was observed (p < 0.01). The geometric mean and median p-value among all significant p-values are also indicated and highlighted in blue. (D) Representative examples of the effects of circadian phase on dwPLI. LSmeans and SEM indicate circadian phase-dependent estimates at 60 degrees (∼ 4 h) bins measured across all studied sleep intervals and EEG derivations presenting a significant circadian modulation as shown in A. Data are double plotted to enhance the visualization of circadian rhythms. The green vertical line marks the dim-light melatonin onset (DLMO; 0° circadian phase), and the grey shaded area represents the average melatonin profile. Type III fixed effects are reported, with only statistically significant p-values (α = 0.01) indicated. Detailed statistical results are provided in Dataset S5.xlsx.

During the spontaneously occurring WAKE periods within the 9h20 min sleep episodes, the dissipation of sleep pressure led to a significant decrease in global FC (PC1), particularly in the theta, sigma, and beta frequency bands (Fig. 7*B* and Dataset S4.xlsx). In the delta and alpha bands, global FC initially declined in the first half of the night but showed a subsequent increase during the final third of the sleep episode. The effects on PC2 were characterized by an increase in topographically distributed FC in the alpha band and a decrease in the beta band, with most other frequency bands showing a non-monotonic shift in FC across the sleep episode similar to PC1 (Fig. 7*E* and Dataset S4.xlsx).

The exploratory topographical analysis revealed that the number of electrode pairs showing a significant elapsed-time-in-sleep-dependent effect during wake was relatively small compared to REM and NREM (Fig 8*A* and Dataset S5.xlsx). The electrode pairs which showed such a significant effect of time in sleep were primarily characterized by a decline in connectivity over time, except for the alpha band, which showed both decreases and increases in connectivity throughout the sleep episode (Fig 8*A*, *B* and Dataset S5.xlsx). The only electrode pair showing a significant effect in the delta band exhibited an initial decline, followed by a return to baseline connectivity levels (Fig 8*B*).

**Fig 7.**
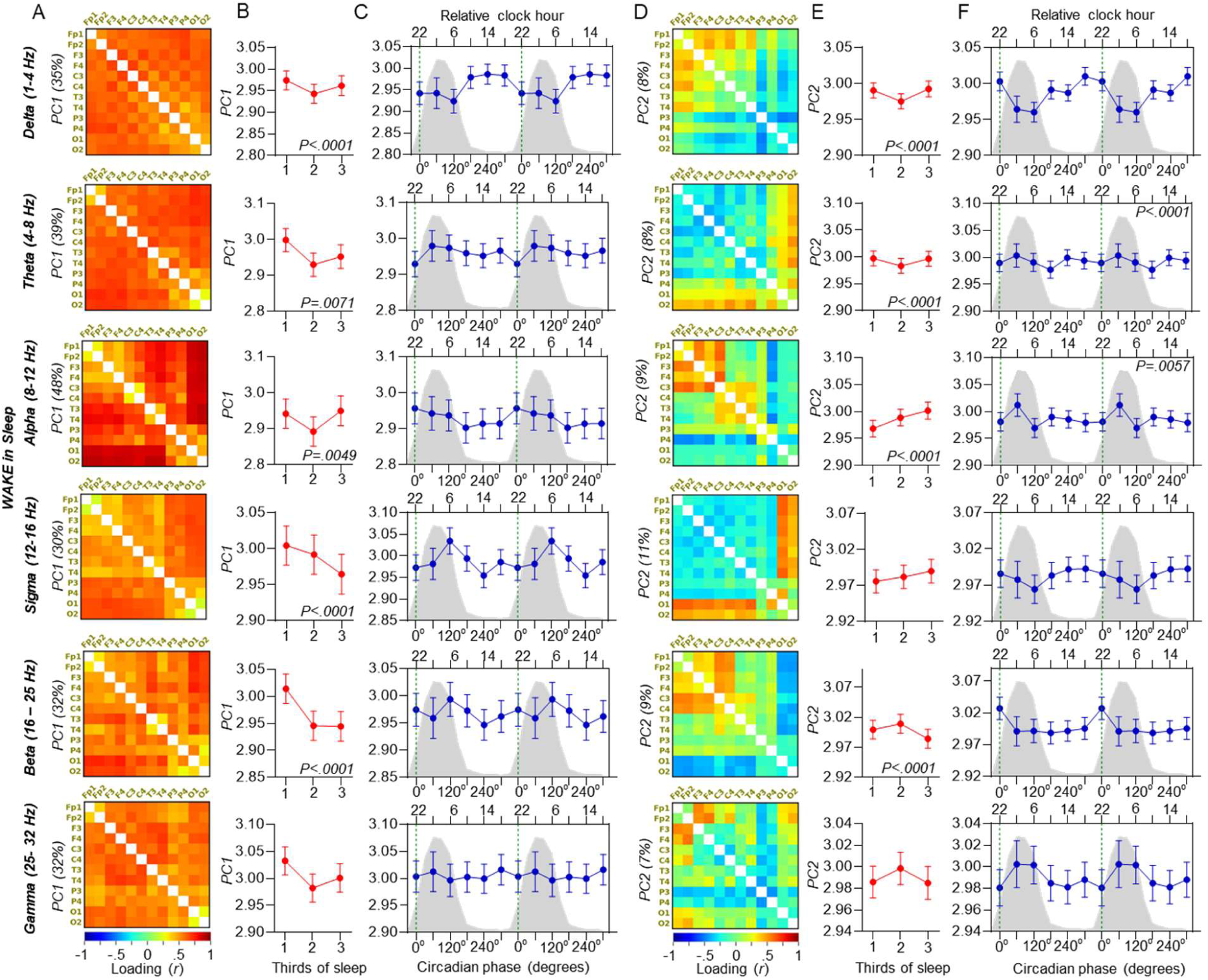
Principal component analysis of dwPLI-based functional connectivity during wakefulness in the time-in-bed period: effects of sleep progression and circadian phase. (A and D) Loading values of individual electrode pairs on PC1 and PC2, for delta, theta, alpha, sigma, beta and gamma frequencies. Values between brackets along the vertical axes represent the variance explained by the PC. (B and E) The modulation of PC1 and PC2 by elapsed time in sleep. Least square means (LSMeans) and standard errors of the mean (SEM) are presented for each 186.7- minute interval (i.e., one-third of the sleep episode), averaged across all studied circadian phases. (C and F) the circadian modulation of PC1 and PC2, with LSMeans and SEM shown for each 60° (∼4-hour) circadian phase bin, averaged across all sleep intervals. Data are double plotted to enhance the visualization of circadian rhythms. The green vertical line marks the dim-light melatonin onset (DLMO; 0° circadian phase), and the grey shaded area represents the average melatonin profile. Type III fixed effects are reported, with only statistically significant p-values (α = 0.01) indicated. Detailed statistical results are provided in Dataset S4.xlsx.

**Fig 8.**
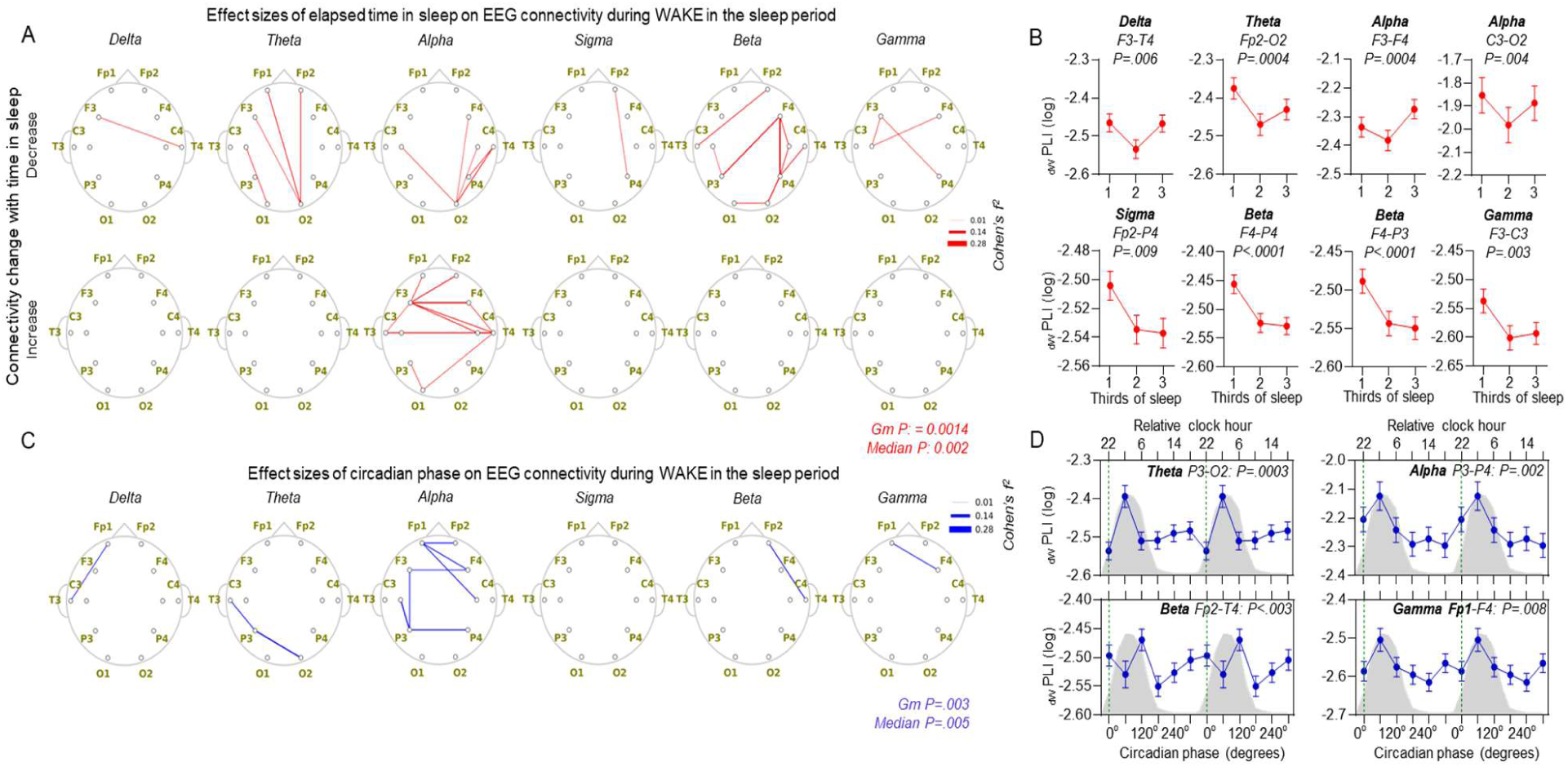
Topographical patterns of functional connectivity (dwPLI) during wakefulness in the time-in-bed period as a function of sleep progression and circadian phase. (A) Red lines represent the magnitude of effect sizes for elapsed time in sleep for electrode pairs, where a significant main effect of this predictor was observed (p < 0.01). Effect sizes for electrode pairs that showed a time-in-sleep–dependent increase or decrease in functional connectivity are shown separately. The geometric mean and median p-value among all significant p-values are also indicated and highlighted in red. (B) Representative examples of the effects of elapsed time in sleep on dwPLI. Least square mean (LSmeans) and standard error of the mean (SEM) are presented indicating sleep-dependent estimates at each 186.7- minute interval (third of the sleep episode) measured across all studied circadian phases. (C) Blue lines represent the magnitude of effect sizes for elapsed time in sleep for electrode pairs, where a significant main effect of this predictor was observed (p < 0.01). The geometric mean and median p-value among all significant p-values are also indicated and highlighted in blue. (D) Representative examples of the effects of circadian phase on dwPLI. LSmeans and SEM indicate circadian phase-dependent estimates at 60 degrees (∼ 4 h) bins measured across all studied sleep intervals and EEG derivations presenting a significant circadian modulation as shown in A. Data are double plotted to enhance the visualization of circadian rhythms. The green vertical line marks the dim-light melatonin onset (DLMO; 0° circadian phase), and the grey shaded area represents the average melatonin profile. Type III fixed effects are reported, with only statistically significant p-values (α = 0.01) indicated. Detailed statistical results are provided in Dataset S5.xlsx.

### Both global and distributed FC are modulated by circadian phase in a topographical, brain-state and frequency-specific manner

The circadian modulation of FC was statistically significant across all brain states and frequency bands with fewer electrode pairs showing a significant modulation of FC compared to the effects of elapsed time in sleep (Fig 3*C* to Fig 8C and Dataset S4.xlsx).

In NREM sleep, global FC (PC1) in the delta band (Fig. 3*C*) and topographically distributed FC (PC2) in the theta band (Fig. 3*F*) showed significant circadian modulation, both peaking during the circadian day, i.e. when plasma melatonin levels are low (Table S5.csv). Exploratory topographical analysis confirmed these effects and further revealed significant circadian modulation for some electrode pairs in all other frequency bands except in beta and gamma (Fig. 4*C* and Dataset S5.xlsx). In the theta band, significant electrode pairs were primarily clustered over centro-posterior cortical regions, whereas in the sigma band, effects were more prominent over fronto-central areas. FC generally peaked during the circadian day, except in the sigma band, where FC was highest during the circadian night, i.e. when plasma melatonin concentrations are high (Figs. 3*C*).

In REM sleep, only global FC (PC1) in the alpha and beta bands showed significant circadian modulation, peaking at the end of the circadian day, i.e. near the onset of the nocturnal surge in melatonin (Fig. 5*C* and Table S5.csv). Exploratory analysis confirmed strong circadian modulation in the alpha band across multiple electrode pairs, peaking in the second half of the circadian day (Figs. 6*C*, D and Dataset S4.xlsx). In contrast, beta and gamma bands exhibited circadian modulation that peaked during the circadian night.

During wakefulness within the sleep episode, circadian modulation of PC1 did not reach significance (*P*>0.01) (Fig. 7*C*), while PC2 showed significant modulation primarily in the theta and alpha bands (Fig. 7*F* and Dataset S4.xlsx). Exploratory topographical analysis supported these findings, revealing a limited number of electrode pairs, mostly in the alpha band, that exhibited significant circadian modulation, all peaking during the circadian night (Figs. 8*C*, D and Dataset S5.xlsx).

In summary, FC during wakefulness tended to peak during the circadian night, whereas in NREM and REM sleep, the circadian phase of peak FC varied by frequency band and principal component / topography.

### Alpha-band-specific global FC during wakefulness increases with time awake and is modulated by the circadian timing of the wake period

We next investigated whether an increase in sleep pressure associated with time elapsed since waking up from a major sleep episode leads to changes in connectivity in wakefulness and whether these changes are opposite to the changes observed when sleep debt dissipates. Wake EEG data collected during 18h and 40 min wake periods starting either in the morning, i.e. at a circadian phase during which humans normally are awake (FdW2), or in the evening (FdW5), as would occur during night shift work (Fig 1*A*), were analysed in a subgroup 12 participants. Wake EEG segments were collected while participants were resting with eyes open during the 2-min long Karolinska Drowsiness Tests (KDT) immediately prior to and after each of six cognitive test sessions scheduled across the wake episode. In total, 365 mins of artifact-free wake EEG segments were analyzed for FC (i.e., wPLI) from 44 electrode pairs (Fig 1*D*). This includes 24 rest EEG wake periods (i.e., KDTs) per person (2 KDTs per cognitive test session per person). Given the limited statistical power inherent in our exploratory, pilot-style analysis of wake-dependent functional connectivity, conducted to elucidate findings from our more extensive sleep data, we took steps to balance Type I and Type II error risks. First, we limited the number of statistical tests by a priori focusing exclusively on the theta and alpha frequency bands, given their well-established associations with both sleep pressure (Finelli, Baumann et al. 2000) and circadian phase (10) and we did not run secondary exploratory analysis. Second, we set our significance threshold at the conventional α = 0.05 to balance sensitivity and reduce the risk of overlooking true effects (i.e., minimize Type II errors).

Despite the smaller dataset, PCA applied to the 45 electrode pairs across the 24 rest EEG wake periods (i.e., KDTs) per person revealed two primary principal components, with interpretations similar to those observed during NREM, REM, and wakefulness within the sleep episode (Figs. S3*A*, *C* and Table S1). Based on the PC loadings PC1 reflected a global FC component (Fig S4*A*), while PC2 represented a topographically more heterogeneous distribute FC component (Fig S4*C* and Dataset S4.xlsx). A linear mixed-effects model including two within-subject factors, time elapsed in wake (thirds of the day: ∼6-hour intervals) and wake period (FdW2 and FdW5), indexing sleep pressure accumulation and circadian phase, respectively, revealed a significant main effect of time elapsed in wake on PC1 in alpha and a significant interaction in both alpha and theta (Fig S4*B* and Table S2). In Alpha global FC showed a steep increase during the 12-hour out-of-phase wake period and followed a non-monotonic pattern during the baseline day. In the theta band, global FC decreased during the baseline wake period, whereas during the 12-hour out-of-phase condition it did not decline and remained overall higher than during baseline (Fig S4*B*). No significant effects were observed on PC2 (Fig S4*D* and Table S2).

### Association of Sleep- and Wake-dependent Functional Connectivity with Cognition

Finally, we examined whether EEG-based functional connectivity (FC) measured during sleep (NREM, REM) and resting wake was associated with cognitive performance, measured in 1,421 cognitive test sessions collected across the forced desynchrony protocol (Fig. 9). To reduce dimensionality, 21 cognitive outcomes were included in the analysis (Fig. 9B), spanning the Karolinska Sleepiness Scale, the Psychomotor Vigilance Test, and multiple N-back tasks (verbal, numerical, pictorial, and integrated at 1-, 2-, and 3-back levels). Two principal components emerged (PC1 = 29% variance; PC2 = 18%) over an eigenvalue of 1, but only PC1 was retained for subsequent analyses due to its robust loadings and interpretability with higher scores indicating higher alertness and working memory accuracy (Fig. 9A–B). Similarly, FC was represented by the PC1 for each brain state and frequency band extracted as described before, only to reduce multiplicity.

**Fig 9.**
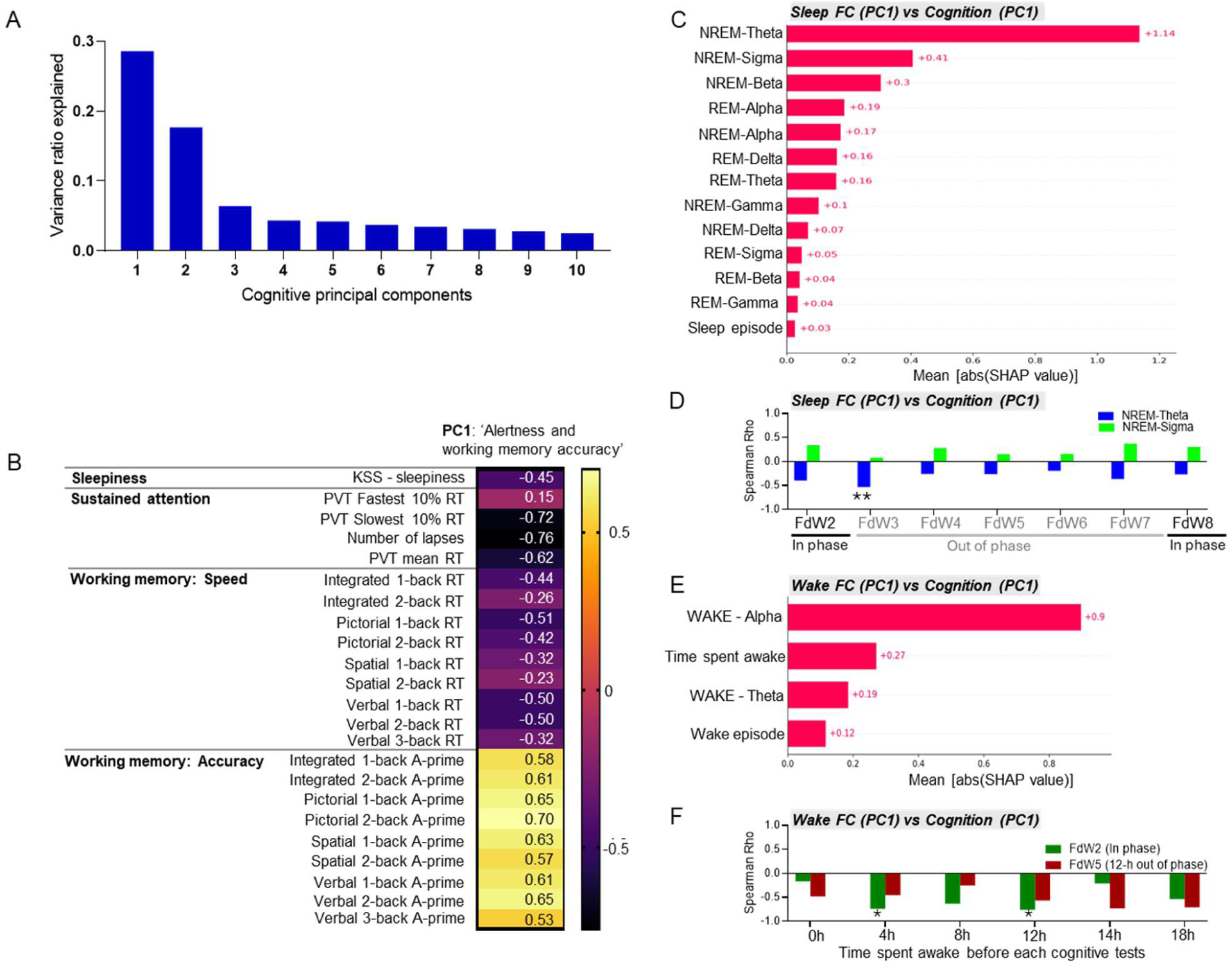
Principal component analysis of cognitive performance and its associations with NREM, REM, and wake functional connectivity. Cognitive performance was measured during the scheduled wake periods. In total, six cognitive test batteries were conducted per wake period. The current PCA included the first cognitive test session following each sleep episode. (A) Explained variance ratio of principal components of cognitive test performance measured across 21 outcome measures covering multiple domains including sleepiness (Karolinska Sleepiness Scale, KSS), vigilance / sustained attention (Performance Vigilance Test, PVT), and working memory (N-back tests). (B) Loading values of individual cognitive outcome measures variables for the PC1’ (‘Higher cognitive alertness and working memory performance’) and PC2 (‘Slower processing but better working memory performance’). Warmer colors indicate stronger positive colder colors stronger negative Pearson correlation coefficients. (C) Mean absolute SHAP values from random forest models assessing the association between functional connectivity (FC) in sleep and cognitive performance (PC1) the subsequent way period across the circadian cycle (N=226). Predictors correspond to the first principal component (PC1) of dwPLI-based connectivity across 66 channel pairs during NREM and REM sleep, calculated separately for each frequency band. Sleep episode (FdN1 to FdN7) was also included in the model. SHAP values represent feature relevance based on game-theoretic Shapley values; their average absolute magnitude reflects each predictor’s global importance in cognitive outcome prediction. (D) Spearman correlations between PC1 of FC in the NREM theta and sigma bands and PC1 of cognitive performance, computed for each sleep episode (N = 34). (E) Mean absolute SHAP values from random forest models assessing the association between functional connectivity (FC) measured in resting wakefulness (Karolinska Drowsiness Test) and cognitive performance (PC1) within the corresponding cognitive test session (N=133). Predictors correspond to the first principal component (PC1) of dwPLI-based connectivity across 45 channel pairs during wakefulness. Additional variables, such as wake period (FdW1 vs FdW5) and time spent awake (test sessions COG1 to COG6), were included in the respective models. (F) Spearman correlations between PC1 of FC in the wake alpha band and PC1 of cognitive performance, computed separately for each cognitive test session and for both in-phase and out-of-phase wake periods (N = 12). Full statistical results are reported in Table S3 to S6.

To investigate the relationship between NREM- and REM-dependent FC (PC1 – ‘global connectivity’) and cognitive performance (PC1 – ‘alertness and working memory accuracy’) on the subsequent day, we followed a two-step procedure. In the exploratory step, we applied both standard multivariate association testing using ordinary least squares (OLS) regression and machine learning-based prediction approaches, including linear regression as well as ridge, lasso, and random forest regressions to identify relevant features (see methods). Despite differences in methodology, both approaches highlighted similar predictors, suggesting convergence between statistical inference and predictive modelling.

The exploratory models included FC PC1 scores across six frequency bands in both NREM and REM (12 predictors), along with the sleep episode (FdN1 to FdN7) to account for repeated structure (N = 34; 226 observations in total) (Fig. 1A). Predictive accuracy ranged from R² = 0.29 (OLS) to –0.17 (lasso), nonetheless, several converging associations appeared. In OLS, NREM-theta FC was negatively related to cognition (β = –0.97, p < 0.00001), while NREM-sigma (β = 0.57, p = 0.001) and NREM-beta (β > 0, p < 0.01) were positively related. These were also retained in the lasso model (Table S3). Random forest models emphasized similar features, with SHAP rankings highlighting NREM-theta and NREM-sigma (Spearman ρ = 0.69, p = 0.0095 vs linear rankings) (Fig. 9*C*). Thus, across methods, global FC in NREM- theta emerged as the most consistent inverse correlate of cognition, although interpretability is limited by the weak predictive fit.

In a confirmatory step, we ran Spearman correlations between the top two FC predictors (NREM-theta and NREM-sigma) and cognitive PC1, separately for each sleep–wake period. This analysis confirmed that global FC in theta was negatively associated with cognitive PC1 while in sigma this association was positive but considerably weaker and non-significant. The associations were further modulated by the circadian timing of sleep–wake episodes (Fig. 9D and Table S4).

We next examined the relationship between wake-dependent FC PC1 scores measured during resting wakefulness in the Karolinska Drowsiness Test (KDT) across six test sessions and two wake periods, habitual wake (FdW2) and 12-h out-of-phase wake episodes (FdW5), and the corresponding cognitive PC1 scores in a smaller pilot sample (N = 12; 133 observations in total). The initial exploratory multivariate linear and non-linear regressions analyses included four predictors represented by wake-dependent FC PC1 (global connectivity) scores in the alpha and theta bands as well as test session (COG1–COG6) and wake period (FDW2, in-phase; FDW5, out-of-phase), given the repeated structure of the data. The dependent variable was represented by the PCA-derived cognitive scores for cognitive PC1 for each individual test session and studied wake period. Despite a poor predictive performance (linear: R² = –0.08; RF: R² = 0.06) both linear regression coefficients and rankings consistently identified WAKE-alpha FC as the most relevant predictor (Fig 9*E*) with higher alpha FC predicting poorer cognitive scores with strong significance (β = – 0.67, *p* < 0.00001, OLS), and was selected by the lasso model (Table S5). Confirmatory Spearman correlation analyses between wake–alpha and cognitive PC1 showed that alpha-band global FC was negatively and significantly associated with cognitive performance, and this association was modulated by test session and the circadian timing of the wake period (Fig 9*F* and Table S6).

In summary NREM-theta connectivity (negative association) and wake-alpha connectivity (negative association) emerged as the most consistent predictors of reduced alertness and working memory efficiency. These findings highlight potential frequency-specific links between FC and cognition.

## Discussion

Separation of sleep-wake and circadian contributions to brain oscillations demonstrates that functional brain connectivity differs across sleep–wake states, is modulated by circadian phase, and changes profoundly with the dissipation of sleep pressure. As sleep progresses, functional connectivity increases during NREM sleep and decreases during REM sleep. Functional connectivity in specific frequency bands during NREM as well as during wakefulness associate with cognitive performance. These novel findings imply that sleep-dependent changes in functional connectivity are related to the recovery of brain function during sleep.

### Functional EEG connectivity across brain states

Although prior studies have reported sleep stage-dependent differences in EEG-measured functional connectivity during nocturnal sleep, this is the first study to assess brain state (i.e. sleep stage) effects while simultaneously controlling for both elapsed time in sleep and circadian phase. In addition, we also controlled for the confounding influence of including short EEG segments in the analyses. As a result, the brain state differences observed here may not align precisely with earlier findings, but it is our view that the current approach provides a more accurate assessment of state specific changes in functional connectivity.

The current approach revealed that wake EEG is characterized by a dominant peak in connectivity within the broader alpha range and that connectivity in the alpha range is lower in NREM sleep and lowest in REM sleep. Wakefulness, however, does not consistently exhibit higher connectivity than other brain states across all frequency bands. For example, connectivity in the sigma band, which primarily reflects sleep spindle activity, is highest in NREM2. Our results support earlier findings indicating that alpha-band functional connectivity is a key feature distinguishing wakefulness from REM sleep and may reflect the degree of disconnection from the external environment (28). Notably, our results, indicate that EEG connectivity in REM sleep is lower than in slow-wave sleep (SWS) across both the broader alpha and sigma bands, with no frequency band showing higher FC in REM compared to all other brain states. These findings were very similar across the five connectivity measures explored here.

One conclusion of the current findings is that rather than considering REM sleep a paradoxical wake’ state, it could be considered the least wake-like brain state and perhaps the deepest sleep stage.

### Changes in functional EEG connectivity with time elapsed in sleep

The sleep-dependent decline in the PSD in the low-frequency range and in particular slow-wave activity (0.75-4.5 Hz) in NREM sleep has for many years been used as a canonical EEG biomarker for sleep homeostasis and the recovery processes occurring during sleep (12). Slow wave indices were subsequently linked to electrophysiological and molecular markers of synaptic strength, which in turn was linked to synaptic downscaling and the hypothesis that synaptic homeostasis is a core function of NREM sleep (1). The current data demonstrate that the dissipation of sleep debt also has large effects on connectivity measures in NREM sleep and across a wide frequency range. Given that connectivity is also underpinned by functional and structural changes in cortical synapses, we expected some similarities between FC and spectral power dynamics. Contrary to our expectations, connectivity in NREM sleep increases with the dissipation of sleep debt, and in all frequency bands, except the alpha band. This is in line with some reports showing an increase in connectivity from the first to the second sleep cycle in young participants. (15). However, other researchers reported that connectivity in NREM2 decreases as nocturnal sleep progresses (29). These previous studies, however, did not control for circadian phase. Furthermore, focusing on NREM2 alone is insufficient to capture the full dynamics of functional connectivity modulation across all stages of NREM sleep. Topographical distribution and differences across canonical frequency bands also need to be considered. Our data indicate that while NREM sleep is generally marked by a global increase in connectivity, particularly in the sigma band, i.e. in the frequency range of sleep spindles, there is also notable topographical variation. Specifically, certain clusters of electrode pairs show sleep-dependent increases or decreases in connectivity, which may represent region-specific specialized reorganization.

Whereas the increase in connectivity in delta and theta bands contrasts with the decreases in spectral power in these bands, the very widespread increase in connectivity in the sigma band parallels the sleep-dependent increase in sigma power (11). A possible explanation for the observed dissociation between the sleep-dependent changes in power spectral density (PSD) and functional connectivity (FC) in NREM lies in their distinct neurophysiological bases. PSD quantifies the local energy of oscillations, though it can be inflated by volume conduction, whereas phase-based FC captures the synchrony of activity across distributed regions within the same frequency band.

The observations that connectivity in REM decreased with the dissipation of sleep pressure while at the same time NREM connectivity increased underlines the brain state specificity of the dissipation of sleep pressure dependent changes in connectivity. This also highlights an important distinction between functional connectivity and power spectral density measures, because for the latter measure the direction of sleep pressure dependent changes in the lower frequency range are in general similar for NREM and REM sleep (11).

### Circadian modulation in connectivity

EEG connectivity in all brain states is influenced by circadian rhythmicity, but the magnitude and direction of this circadian effect is dependent on brain state and brain topography. This is consistent with previous findings showing circadian modulation of EEG measures, including slow-wave activity and sleep spindle activity in NREM sleep, alpha activity during REM and wakefulness, and markers of synaptic strength such as the slope of slow waves in NREM sleep, particularly in similar forced-desynchrony protocols (10, 11, 25, 30, 31). Although the extent and phase of circadian modulation of these EEG measures depend on brain state, frequency, and topography, our results show that the dominant circadian rhythms of functional connectivity in wakefulness and NREM sleep are approximately 12 hours out of phase.

Specifically, connectivity during wake peaks at the circadian night, whereas connectivity during sleep peaks at the circadian day.

### Functional connectivity and cognitive performance

While the predictive power was weak and the findings should be interpreted with caution, FC during sleep— especially NREM—and during the major wake period showed frequency-specific associations with cognition. This is not surprising as the human brain operates as a network of functionally interconnected regions, the connectivity of which is believed to underpin behavior, cognition, and mood states (32).

However, our findings suggest that the relationship between network synchrony, as measured by EEG, and cognitive performance is a non-trivial one. During NREM sleep, lower global connectivity in the theta band and higher global connectivity in the sigma band (FC–PC1) were associated with better subsequent alertness and working memory performance (cognitive PC1), independent of connectivity in other frequency bands during either NREM or REM sleep. While the increased sigma coupling may reflect more effective thalamocortical network interactions, the link between lower theta connectivity and waking cognitive performance remains unclear.

Notably, previous work has shown that EEG-based functional connectivity in the theta and beta frequency ranges increases with age, whereas connectivity in the sigma range decreases (Ujma et al., 2019). Given that cognitive function normally declines with age, increased theta-band global connectivity may be considered a negative predictor of waking cognitive function, potentially an even stronger one than sigma-based connectivity. This interpretation also aligns with findings in older patients with obstructive sleep apnea where, higher relative frontal theta power during NREM stage 2 was negatively associated with Mini-Mental State Examination scores and was elevated in those with mild cognitive impairment (Chen et al., 2023). These findings suggest that the cognitive benefits of sleep arise from frequency-specific alterations in oscillatory coupling, with distinct roles for sigma- and theta-band synchrony.

In contrast, during wakefulness, higher global alpha connectivity was associated with poorer performance, consistent with previous reports linking excessive alpha synchrony to reduced neural efficiency and impaired cognitive flexibility in both healthy aging and clinical populations (Klimesch, 2012; Babiloni et al., 2016). Resting-state alpha rhythms are generally slowed in mild cognitive impairment, which has been associated with cognitive decline (33). Our observation that global alpha connectivity decreases across sleep, but rebounds with extended wakefulness, may therefore reflect an oscillatory signature of sleep-dependent recovery versus wake-related deterioration of neural efficiency. Alternatively, some associations may reflect stable, trait-like relationships between the brain’s functional neuroarchitecture and cognition (Touroutoglou, Andreano et al. 2015; Seitzman, Gratton et al. 2019).

### Speculative interpretation of the results – Connectivity and Criticality of brain states

Previous research (34, 35) suggests that optimal FC (and thus optimal brain performance) is achieved when the brain operates near a critical state. This critical state supports a balance between flexibility and stability, essential for cognitive processes and adaptability. Perturbations from this critical state, whether due to anesthesia, sleep deprivation, or disorders of consciousness, result in less efficient connectivity patterns. During wakefulness, connectivity levels may increase due to synaptic potentiation, leading to disrupted (super)critical dynamics and less efficient states, whereas sleep acts to restore critical dynamics(35). We demonstrate that the sleep-dependent recovery process is characterized by a pronounced reduction in waking-state connectivity, particularly in the alpha band, which has been linked to waking vigilance in both our and previous research (10). And indeed, alpha band emerges as a unique marker. Not only was alpha FC the only frequency to show a systematic decrease over the course of both NREM and REM sleep, but daytime wake EEG also revealed that lower global alpha connectivity strongly predicted better subsequent cognitive performance. This robust negative relationship suggests that efficient network desynchronization in the alpha band, both during sleep and wake, may index a neural state optimized for vigilance and information processing.

Although sleep recovery exerts broad, global effects on synaptic strengths, as described by the synaptic homeostasis theory, the stage-specific patterns we observe (rising connectivity in NREM and declining connectivity in REM) suggest that the renormalization of large-scale network interactions and information-processing capacity depends on a coordinated, multi-phase recovery process rather than a uniform downscaling. That is the coordinated alternation between NREM- and REM-mediated network recalibration with opposite directions may be critical for rebalancing neural integration and information-processing capacity. This aligns with theory suggesting that the inhibitory, collothalamic mechanisms of NREM sleep, in nightly tandem with the excitatory, lemnothalamic mechanisms of REM sleep, support the consolidation of new memories and their interleaved reconsolidation with existing memory representations (36).

### Limitations and Strengths of the Current Study

The results presented here are based on a rigorous design and a large data set; nevertheless, the study has several limitations. High-density EEG and source analysis which would have allowed a more comprehensive connectivity analysis was not implemented. As such, our EEG connectivity analyses cannot directly be linked to distinct functional brain connectivity networks. Our experimental manipulation was limited to sleep-wake timing. We did not perturb physiological brain activity and connectivity via electric or magnetic stimulation.

Even though our EEG methodology may be limited, the data reveal aspects of the sleep process which have not previously been reported. The finding presents a unique demonstration of the separate contribution of the sleep-wake cycle and circadian rhythmicity to FC in wake, NREM and REM sleep. Previous reports were based on small sample sizes, did not separate circadian and sleep-wake dependent contributions and investigated effects of time awake by sleep deprivation which is a challenge to the brain well beyond the challenges experienced during a normal sleep-wake cycle. Previous studies have generally emphasized differences between brain states (e.g., wake vs. NREM) or between sleep stages (e.g., NREM2 vs. SWS). In this large dataset, with repeated measures of both EEG and cognitive performance, we focus on changes in connectivity within brain states rather than differences between them. We have done so because, in our view, understanding how sleep contributes to recovery of brain function requires a description of the dynamics of FC during the sleep episode, i.e. analyse how it changes from the beginning to the end of sleep. Finally, our approach is unbiased because we considered many different frequency bands rather than focusing on only one or two frequency ranges.

### Concluding Remarks

Together, these findings suggest that when the sleep–wake cycle is aligned with circadian rhythmicity, the interaction between sleep homeostasis and circadian processes supports the maintenance of optimal brain connectivity. This balance may be disrupted during misalignment, which could contribute to cognitive performance deficits and neurological conditions associated with disturbances of sleep and circadian rhythms.

For many decades, sleep-wake and circadian aspects of brain function were quantified by sleep staging and simple EEG analysis methods that are dependent on amplitude characteristics of the EEG. More recently amplitude independent measures, such as coherence, were introduced. However, these measures are confounded by volume conduction. These approaches revealed canonical aspects of the sleep process, although it remained to the larger extent unclear how these aspects of the EEG and sleep process associate with brain function. Novel approaches to EEG analysis, such as those used here, combined with comprehensive assessment of brain function, implemented in protocols that separate the contribution of sleep and circadian rhythmicity may provide new insights into the process by which sleep and circadian rhythmicity interact to maintain homeostasis of brain function.

## Materials and Methods

The study was approved by the University of Surrey Ethics Committee and conducted in accordance with the Declaration of Helsinki. Written informed consent was obtained from all participants prior to their enrolment.

### Study protocol

The circadian and sleep-wake time-dependent modulation of EEG oscillatory was assessed using a 10-day forced desynchrony (FD) protocol, conducted in the Surrey Sleep Research Centre at the University of Surrey. The protocol and associated procedures have been described in detail elsewhere (25, 30, 31). Briefly, following a baseline day (FdW1) and night (FdN1), participants were scheduled to a 28-h sleep-wake cycle comprising 18 h and 40 min of wakefulness in dim light (< 5 lx) and 9 and 20 min of sleep opportunity in darkness (Fig 1a). Throughout the 10-day period, participants resided in the clinical research facility and followed a standardized routine including scheduled mealtimes and cognitive testing sessions without access to information about clock time. Participants were continuously monitored during wake periods by a member of staff to ensure wakefulness and followed a strict schedule without physical exercise. During the scheduled wake periods, participants spent most of their time in a lounge where they were allowed to chat with each other or with a member of staff, listen to music, and watch movies and TV series from a pre-selected list.

### The assessment of the circadian phase

Blood samples were collected hourly during the 1st, 4th, and 7th 28-h forced desynchrony sleep-wake cycles (FdW1, FdW4, and FdW7) for determination of melatonin concentrations and assessment of circadian phase (25, 30, 31) (Fig 1a). Blood melatonin concentration was quantified using radioimmunoassay (Stockgrand Ltd, Guildford, UK). The time corresponding to the 25% of the daily melatonin amplitude range [dim light melatonin onset (DLMO)] was assigned circadian phase zero (37, 38). Circadian period was derived from the linear regression fitted to the 3 DLMOs measured at FD1, FD4, and FD7 for each participant (τ = 24 h + slope) (39).

### Assignment of circadian phase and elapsed time in sleep and time in wake period to dependent variables

To estimate the independent contribution of circadian phase and time since the start of the sleep and wake episodes, to the outcome measures, we grouped the continuously recorded EEG data into time bins and assigned to each of those time bins a circadian phase and the sleep-wake dependent index as described earlier (25, 30).

PSD and connectivity measures were computed for all consecutive 20-minute intervals from the start to the end of each sleep episode. We then assigned a circadian phase and a time since the start of the sleep episode to each of the 20-minute intervals. For this, we used the 9 h 20 min sleep episodes across FdN1 to FdN7. In the next step, data from the sleep and wake epochs (e.g., wake, NREM, REM) and the consecutive 20-minute intervals (spectral data and connectivity measures) were averaged in 6 circadian phase bins, each of 60° (∼ 4-hourly bins), and three sleep-dependent time bins (around 186.7 mins each) for each sleep episode.

For the wake-duration-dependent EEG analysis, we used the resting state wake EEG data collected during the Karolinska Drowsiness Tests (KDT) with eyes open and fixating a point for two minutes. These KDT sessions were scheduled immediately prior to and following each of the six cognitive test sessions during the 2^nd^ and the 5^th^ wake period. The 2^nd^ wake period started at habitual wake time (e.g. approximately 8am) and the 5^th^ wake period started 12 hours later (Fig 1). Similar to the sleep-duration-dependent analyses, the wake data were averaged per third of the 18h-40 min wake periods.

### Assessment of cognition

Participants completed approximately 40 minutes of computerized cognitive testing at three-hour intervals. For the present analyses, we examined subjective sleepiness using the Karolinska Sleepiness Scale (KSS; administered at the beginning and/or end of each session), objective vigilance and sustained attention via the 10-minute Psychomotor Vigilance Test (PVT), and working memory and executive function using 1-, 2-, and 3-back tasks. A detailed description of these measures has been published elsewhere(25, 40). The KSS is a 9-point Likert scale ranging from 1 (extremely alert) to 9 (very sleepy), with higher scores indicating higher levels of subjective sleepiness. From the PVT, we derived four outcome measures: mean reaction time (RT), number of lapses (RT > 500 ms), and the inverse of the slowest and fastest 10 % of RTs. For the n-back tasks, performed in four variants (integrated, pictorial, spatial, and verbal), we extracted two main outcome measures for each variant: RT and A′ (A-prime). A′ is a nonparametric sensitivity index that quantifies a participant’s ability to discriminate targets (hits) from non-targets (false alarms), ranging from 0.5 (chance performance) to 1.0 (perfect discrimination).

### Polysomnographic (PSG) and wake EEG assessment

EEG was recorded throughout the forced desynchrony protocol during both sleep episodes and repeated cognitive test sessions during the wake episodes. For the current analyses, we analyzed the complete EEG data acquired during 231 sleep episodes from 34 participants (18 females, age: 25.1 ± 3.4). All participants presented a healthy sleep profile based on their baseline adaptation night which also served as a clinical sleep screening with a full clinical EEG-PSG setup (12 EEG channels, EOG, EMG, thoracic belt, a nasal airflow sensor, a microphone, and leg electrodes). During the FD protocol, sleep and wakefulness were monitored by basic polygraphy (EMG, ECG, and EOG) with extended monopolar EEG montage which covered the major brain areas (Fp1, Fp2, F3, F4, C3, C4, T3, T4, P3, P4, O1, and O2) following the international 10–20 system. The ground and common reference electrodes were placed at FPz and Pz, respectively. Two referencing schemes were applied. For power spectral density (PSD) analysis, EEG derivations were re-referenced offline to the contralateral mastoid (A1 or A2). For connectivity analyses, a common reference was used across all connectivity metrics. Polysomnographic (PSG) data were recorded using Siesta 802 amplifiers (Compumedics, Abbotsford, Victoria, Australia).

During the scheduled wake periods, EEG data were collected using the TEMEC Vitaport 3 system (TEMEC Instruments B.V., Kerkrade, The Netherlands). The EEG montage was similar to the montage used for the sleep EEG recording except that Fp1 and Fp2 channels were not used due to eye blinking artifacts. For the current analysis, wake EEG from twelve participants out of 34 were included in the analysis (nine men, and three women). This is because the paper focuses on the sleep data, while the wake EEG analyses represent a pilot investigation. The participants whose EEG data were analyzed did not differ from the rest of the sample in terms of demographic characteristics or baseline sleep parameters.

EEG data were stored at 256 Hz. The low-pass filter was set at 70 Hz and the high-pass filter was set at 0.3 Hz. Electrode impedance was kept below 5 kΩ. As reported earlier (30) sleep staging was performed in 30 s epochs according to the Rechtschaffen and Kales criteria (41) by one experienced sleep researcher (ASL) whose scoring showed a concordance exceeding 90% when compared to a standard scored data set.

### EEG power spectrum and connectivity analyses

All artifact-free EEG segments of the sleep stages of interest (WAKE, NREM1, NREM2, SWS, REM) were concatenated within consecutive 20 min time intervals between lights out and lights on spanning 9 hours and 20 minutes (28 intervals) sleep episode. In total, 76,264 min (1771 hrs) artifact-free EEG sections were analyzed for the sleep episodes including 58,155 min for NREM and 18,109 min for REM as well as 4865 min for WAKE in the sleep episode and 365 min in the KDTs during the wake episodes with some modulations across the circadian cycle (Fig 1C and D). From the resting state wake EEG data, we processed a two-minute-long EEG segment recorded immediately before and a similar interval recorded immediately after each cognitive test battery.

We chose spectral methods to enable direct examination of frequency-specific effects, allowing us to assess how the circadian-homeostatic interaction modulates functional connectivity in each canonical EEG band. While non-spectral methods combined with bandpass filtering can also provide frequency-targeted estimates, spectral methods offer native, direct frequency resolution, superior phase accuracy, and methodological clarity (42).

Values were either analyzed on a 0.5 Hz bin basis to investigate spectral profiles between 0.5 to 32 Hz or averaged over commonly used frequency bands: delta (0.5-4 Hz), theta (4-8 Hz), alpha (8-12 Hz), sigma (12-16Hz), beta (16–25), and gamma (25 - 32 Hz). All measures were computed across standard EEG frequency bands (delta, theta, alpha, sigma, beta, gamma) and separately for each brain state.

Computations used Python’s NumPy and SciPy packages (43, 44).

Detailed descriptions of the Power Spectral Density and Local Connectivity measures are provided in the Supplementary Appendix.

### Evaluation and selection of functional EEG connectivity metrics

While we ran the initial analysis characterizing the spectral profile of FC across different brain states using five different connectivity measures, we chose to run the in-depth analysis using dwPLI. The choice of dwPLI as the primary connectivity metric is well-supported by its correlation profile. It exhibits a very high Pearson correlation with wPLI (0.95) and strong correlations with PLI (0.81) and ImCoh (0.66), suggesting that it effectively captures the core aspects of phase-based connectivity shared across these widely used measures. At the same time, its low correlation with phase coherence (0.10) confirms its resistance to volume conduction and common source artifacts, which often inflate coherence-based estimates. This balance between specificity and robustness makes dwPLI a particularly suitable choice as it summarizes the essential structure of inter-regional interactions without being compromised by methodological limitations inherent in simpler or more artifact-prone metrics.

The selection of dwPLI was further supported by superior fit statistics for the mixed-effects model, specifically lower Akaike Information Criterion (AIC) and Bayesian Information Criterion (BIC) values, compared to those obtained using ImCoh.

### Principal component analysis of functional brain connectivity and cognition

We applied Principal Component Analysis (PCA) to the large, multivariate datasets related to FC and cognition including all electrode pairs and sleep/wake episodes to reduce dimensionality and investigate their underlying structure and relationships. For the FC data PCAs were run separately for each brain state and frequency band. The employed imputation strategy for handling missing values in the datasets consisted of removing all rows containing missing data. All columns were standardized to ensure that all variables contributed equally to the analysis. In this instance, standardization was applied by dividing each feature by its standard deviation. PCA was then conducted using a specified number of principal components; in this case, 10 components were chosen to represent the underlying structure of the data.

After fitting the PCA model, we extracted the component loadings and scores and calculated the explained-variance ratio for each component to quantify the proportion of total variance accounted for. Across all EEG connectivity matrices, PC 1 captured the majority of meaningful variance and was marked by a distinct elbow in the scree plot. In a few frequency bands during NREM and wakefulness, PC2 also accounted for a relatively large share of variance (>10%). All remaining components each explained less than 5 % of the variance. In line with common PCA practices (e.g., Kaiser’s criterion and the scree test) and to maintain consistency across brain states and frequency bands, we therefore retained only the first two components for all subsequent analyses.

To examine the association between FC and cognitive performance, a principal component analysis (PCA) was first conducted across all cognitive test sessions, in line with the approach used for FC. In the first analysis, which focused on the association between sleep-dependent FC and cognitive PC1. The cognitive PC scores were analyzed as averaged for the entire wake episode (i.e., 6 test sessions) following each sleep episode (see Fig. 1A). In contrast, the second pilot analysis, targeting the association between FC measured during habitual (FdW2) and out-of-phase (FdW5) wakefulness episodes (Fig. 1A) and cognition, included both FC and cognitive PC scores from all six cognitive assessments conducted within each wake episode. Only the first principal component from the cognitive PCA was retained, reducing dimensionality while preserving the main axes of variance relevant to our outcome.

### Statistical analysis

All analyses were conducted in SAS 9.4 and visualized using GraphPad Prism 9.

### Power and Connectivity spectrum analysis

In the first analysis, the dependent variables were spectral power and electrode-pair connectivity, measured with five metrics (coherence [COH], imaginary coherence [ImCOH], phase-lag index [PLI], weighted PLI [wPLI], and debiased wPLI [dwPLI]), computed at 1 Hz resolution across each sleep stage (NREM1, NREM2, slow-wave sleep, REM, and wake) during the scheduled sleep episodes. The PSD and connectivity outcome measures were averaged across all EEG channels and channel pairs, respectively, as topography was not evaluated at this stage. To examine the influence of circadian phase, elapsed time in sleep, and sleep stage on these dependent measures we conducted linear mixed-effects modelling for each log transformed PSD and connectivity outcome measure across the frequency spectrum and across all 231 scheduled sleep episodes. The fixed effects included three within-subject factors: **circadian phase** (six levels, using 60° bins), **elapsed time in sleep** (divided into three approximately 3.1-hour intervals), and **sleep stage** (Wake, NREM1, NREM2, NREM3, and REM). To accommodate the repeated measures structure of the data, we created a composite factor representing each unique combination of these three conditions and treated this as the unit of repetition nested within each participant and session. Random intercepts were included to account for inter-individual differences, and a compound symmetry covariance structure was specified to model within-subject correlations across repeated measurement occasions. To improve the accuracy of standard error estimates, we used the Kenward–Roger method for degrees of freedom estimation. The analysis focused on estimating and testing the main effects of each factor on connectivity as well as the two-way interactions of factors *sleep stage* with *circadian phase* and *elapsed time in sleep*. To control for Type I error, the significance threshold was adjusted to α = 0.01 (see section on Correction for Multiplicity).

### Topographical analysis of EEG connectivity

The topographical analysis comprised two consecutive steps. In the primary analysis, the dependent variables were the principal component (PC) scores derived from a PCA on dwPLI-based electrode-pair connectivity. Connectivity was calculated within six canonical frequency bands (delta, theta, alpha, sigma, beta, and gamma) across three consolidated brain states (NREM, REM, and Wake) during 231 sleep episodes, and during baseline wakefulness (FdW2) and 12 h out-of-phase wakefulness (FdW5) in a subset of participants. Although the PCA reduced the 66 individual electrode-pair connectivity values to a smaller set of components, the original PC loadings enabled topographical interpretation of each component.

For the sleep episodes, we focused on the first two PCs, three brain states, and six frequency bands, yielding 2 × 3 × 6 = 36 unique models. Each model examined a single PC, brain state and frequency-band combination. To control for Type I error, the significance threshold was adjusted to α = 0.01.

For the wake EEG data (measured during the two wake episodes), we again used the first two PCs but restricted our analysis to the theta and alpha bands, given their established roles in homeostatic and circadian modulation, resulting in 2 × 2 = 4 models, each examining a single PC–frequency-band combination. To balance statistical rigor with our small exploratory sample (N = 12) due the intensive manual artifact rejection required, we limited the number of tests and set the significance threshold at α = 0.05.

In the second step, we conducted a more exploratory topographical analysis by examining dwPLI-measured connectivity for each electrode pair individually. In both the primary and the secondary analyses, we applied the same linear mixed-effects modelling approach across brain states and canonical frequency bands. The fixed effects included two within-subject factors: circadian phase (six levels, using 60° bins) and time elapsed in sleep (divided into three approximately 3.1-hour intervals).

For the sleep data, comprising 66 EEG channel pairs, three brain states (NREM, REM, and Wake), and six frequency bands, this resulted in 66 × 3 × 6 = 1 188 unique models. For the wake EEG data during the two wake episodes, comprising 45 channel pairs and two frequency bands, we tested 45 × 2 = 90 models. Each model analysed a single channel–state–band combination.

Since each subject contributed up to seven sleep episodes (SEs), and within each SE up to three sleep intervals (i.e., thirds of the night), we specified a compound-symmetry (CS) covariance structure on time elapsed in sleep nested within circadian phase (i.e., repeated ElapsedTimeInSleep (CircadianPhase) / subject = Subject*SE type = CS) to parsimoniously model within-episode correlations.

For the analysis of connectivity during the two wake episode, the two within-subject factors were elapsed time in wake (divided into thirds of each wake period) and wake episode (in-phase vs. out-of-phase), including their two-way interaction. Mixed models included a random intercept for each participant to account for between-subject variability. Model degrees of freedom were adjusted using the Kenward–Roger method (ddfm = kr) to improve the accuracy of standard errors and test statistics.

### Association analysis between Connectivity and Cognition

All multivariate analyses were conducted in Python (v3.8) (45) using standard scientific computing libraries, including numpy, scipy, pandas, scikit-learn, statsmodels, and shap (46–48). Functional connectivity was calculated using the debiased weighted Phase Lag Index (dwPLI), separately for NREM, REM, and resting wakefulness. For each frequency band and sleep–wake stage, 66 (sleep) or 45 (wake) channel-pair values were reduced via principal component analysis (PCA) to a single latent feature: the first principal component (PC1), representing global connectivity for that condition. Cognitive performance was likewise reduced through PCA across 21 behavioral outcomes spanning sleepiness ratings, vigilance, and working memory tasks. The first cognitive component (PC1), interpreted as a general alertness and working memory factor, served as the target variable. For sleep analyses, PC1 scores were averaged across the six behavioral sessions following each sleep episode.

To examine the relationship between functional connectivity and cognition, a two-step analysis was applied. In the exploratory phase, both association and prediction models were employed. Multivariate association was assessed using Ordinary Least Squares (OLS) regression via the statsmodels package, yielding estimated regression coefficients, p-values, and 95% confidence intervals. In parallel, prediction was performed using linear regression, ridge regression with cross-validated regularization strength (49), lasso regression (also with CV-based regularization) (50), and random forest regression with 100 estimators (51). Feature values were standardized before modeling. The outcome variable was negated prior to training to align the direction of prediction across datasets. Subject identity was optionally included as a predictor to evaluate the extent to which individual baselines influenced model performance.

To avoid data leakage, an 80/20 train/test split was used, ensuring that data from a given participant appeared only in one of the sets. Train/test splits were group-aware and stratified by subject identity. Missing values were handled via mean imputation. To interpret feature relevance in tree-based models, SHAP values were computed using the shap package. SHAP (SHapley Additive exPlanations) is a method for quantifying the contribution of each feature to model output, based on cooperative game theory, and is conceptually related to feature importance measures but offers consistent, model-agnostic attributions (52–54).

In the confirmatory phase, the most relevant predictors identified during the exploratory phase were tested using univariate Spearman correlations with cognitive PC1 scores, calculated separately for each sleep– wake episode. Spearman correlations were performed in SAS 9.4.

### Output and Diagnostic Tracking

For each model, we extracted key analytical outcomes, including the estimated least-squares means for the levels of both predictors, Type III tests of fixed effects (F-values and associated p-values), and standard fit statistics such as AIC and BIC. We also recorded the residual skewness of the fitted model and documented whether the final model used a log-transformed or untransformed outcome variable. To visualize statistical effect sizes, we calculated Cohen’s *f^2^* effect size (55):

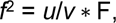

where *u* and *v* are, respectively, the numerator and denominator degrees of freedom of the F statistic used to determine the corresponding main or interaction effect in the general linear mixed model analysis.

In the model-selection process, we evaluated the residual-distribution diagnostics, checked for convergence issues (e.g., infinite likelihood), and compared fit statistics, namely the Akaike Information Criterion (AIC) and the Bayesian Information Criterion (BIC). The model exhibiting the lower AIC and BIC was retained as the final model. Correlational analysis was based on Spearman correlations.

### Correction for Multiplicity

We conducted a large number of statistical tests across several multivariate models. Although these tests were not entirely independent, we addressed the risk of Type I error through multiple strategies. First, we set a more stringent significance threshold at α = 0.01. Second, in our primary topographical analysis, we further limited the number of comparisons by reducing dimensionality via a PCA and only the PCs derived from the dwPLI connectivity data were tested. Third, we then carried out an exploratory channel-pair analysis to confirm the primary results, and throughout both stages we reported P values alongside standardized effect sizes, anchoring our main conclusions on the largest and most statistically robust effects.

To contextualize our chosen α = 0.01 relative to a conventional multiplicity correction, we also applied the Benjamini–Hochberg procedure (FDR = 5%). Across all 5100 individual p-values, the largest p-value passing the Benjamini–Hochberg critical threshold was 0.021. Thus, under the Benjamini–Hochberg procedure, tests with p < 0.021 would be deemed significant. Our choice of α = 0.01 is therefore more conservative than controlling the false discovery rate at 5%.

For the wake-episode data, given the smaller sample size and the need to balance Type I and Type II error risks, we reduced the number of analyses and set the significance threshold at the conventional α = 0.05.

## Acknowledgments

This research was supported by a Biotechnology and Biological Sciences Research Council grant (BB/F022883; PI: DJD). DJD is supported by the UK Dementia Research Institute core awards UKDRI- 7005 and CF2023\7 and award UKDRI-7206 , through UK DRI Ltd, principally funded by the UK Medical Research Council; and by the National Institute for Health Research (NIHR) Oxford Health Biomedical Research Centre (BRC), (NIHR203316) . ASL received funding from the Wellcome trust (207799/Z/17/Z) and UKRI (ES/W006367/1) and NIHR (NIHR206949). We thank the staff of the Surrey Clinical Research Center for their help with recruitment, screening and clinical conduct of the study. Drs Ana Slak, Sibah Hasan for their help with data acquisition.

