## Supplementary Appendix for "Circadian and Sleep-Wake Modulation of Functional Connectivity Across Brain Oscillations and States Linked to Cognition in Humans"

### Supplementary Figures

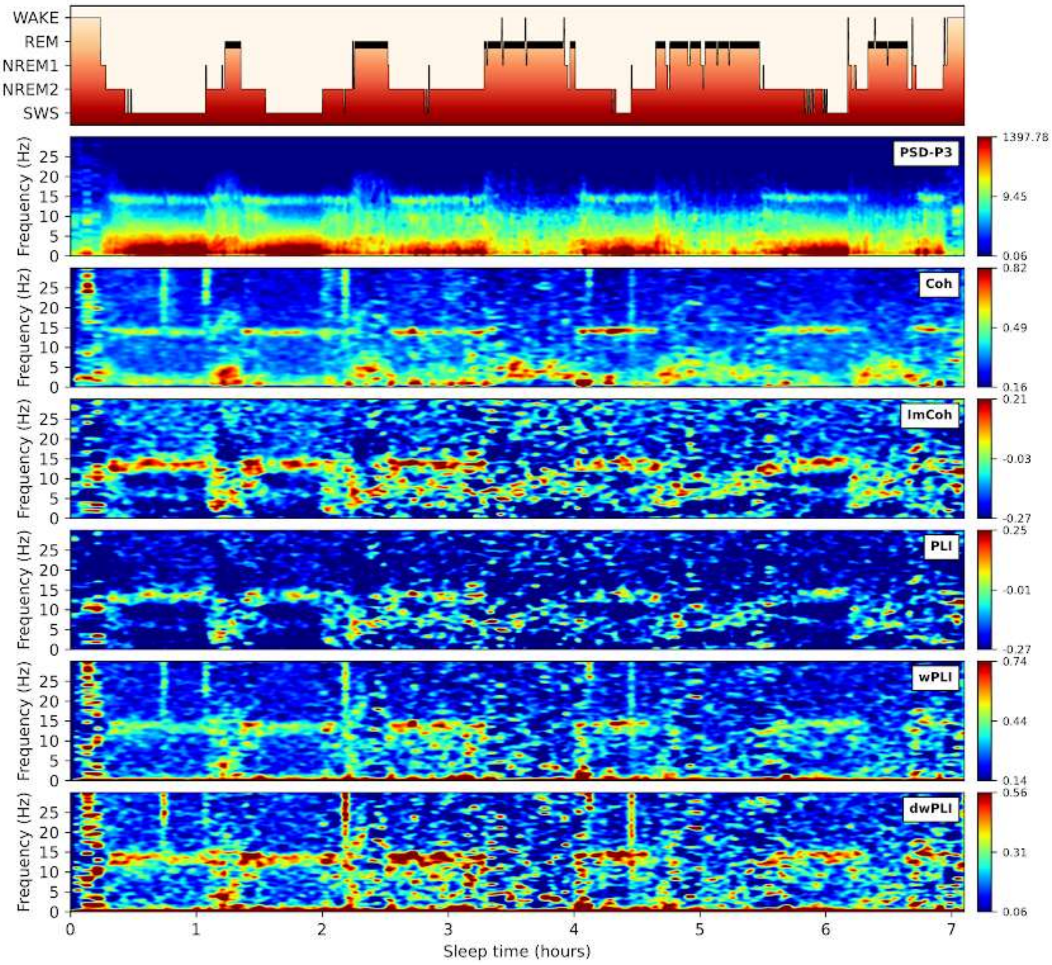


**Fig. S1.** Time course of sleep stages (top panel), power spectral density (PSD) (second panel), and functional connectivity measures (below) including coherence (Coh), imaginary coherence (ImCoh), phase lag index (PLI), weighted PLI (WPLI) and debiased wPLI (DWPLI) measured during a baseline sleep episode in one participant. Warmer colors represent higher spectral power (logarithmic scale) and functional connectivity (FC). Temporal resolution is 10 seconds/pixel, each representing the average of the short-time Fourier transforms of four segments of 4-s long, Hanning tapered windows with 50% overlap. Power spectral density (PSD) values were obtained from channel P3 whereas connectivity values were averaged over all 66 electrode pairs of the 12 channels (FP1, Fp2, F3, F4, C3, C4, T3, T4, P3, P4, O1, and O2).

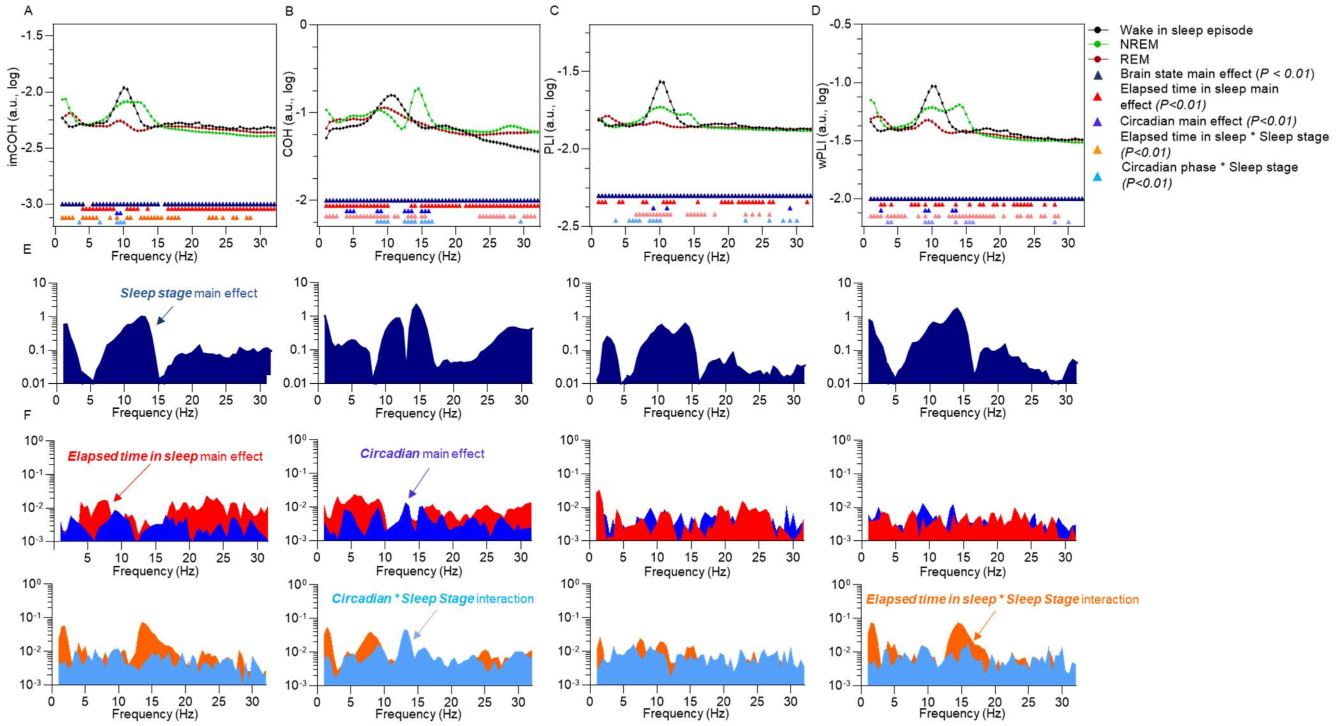


**Fig. S2. Functional connectivity spectra across brain states and all sleep episodes as measured by multiple connectivity metric.** The main effect of vigilance states on (A) Imaginary coherence (ImCOH), (B) Phase coherence (COH), (C) Phase lag index and (PLI) weighted Phase Lag Index (wPLI) across all sleep episodes (231 nights) and averaged across EEG channels and cannels pairs, respectively. The upper panels represent least-square means (Lsmeans) and standard error of the mean (SEM). (E) Effect size (*Cohen's* $f^2$) of main effects of brain state on all connectivity metrics. (F) Effect size of main effects of Elapsed time in sleep (red) and Circadian phase (blue) on all connectivity metrics. Colored triangles in (A) and (B) indicate significant (P<.01) effects. Detailed statistical results are in Dataset S1.xlsx**.**

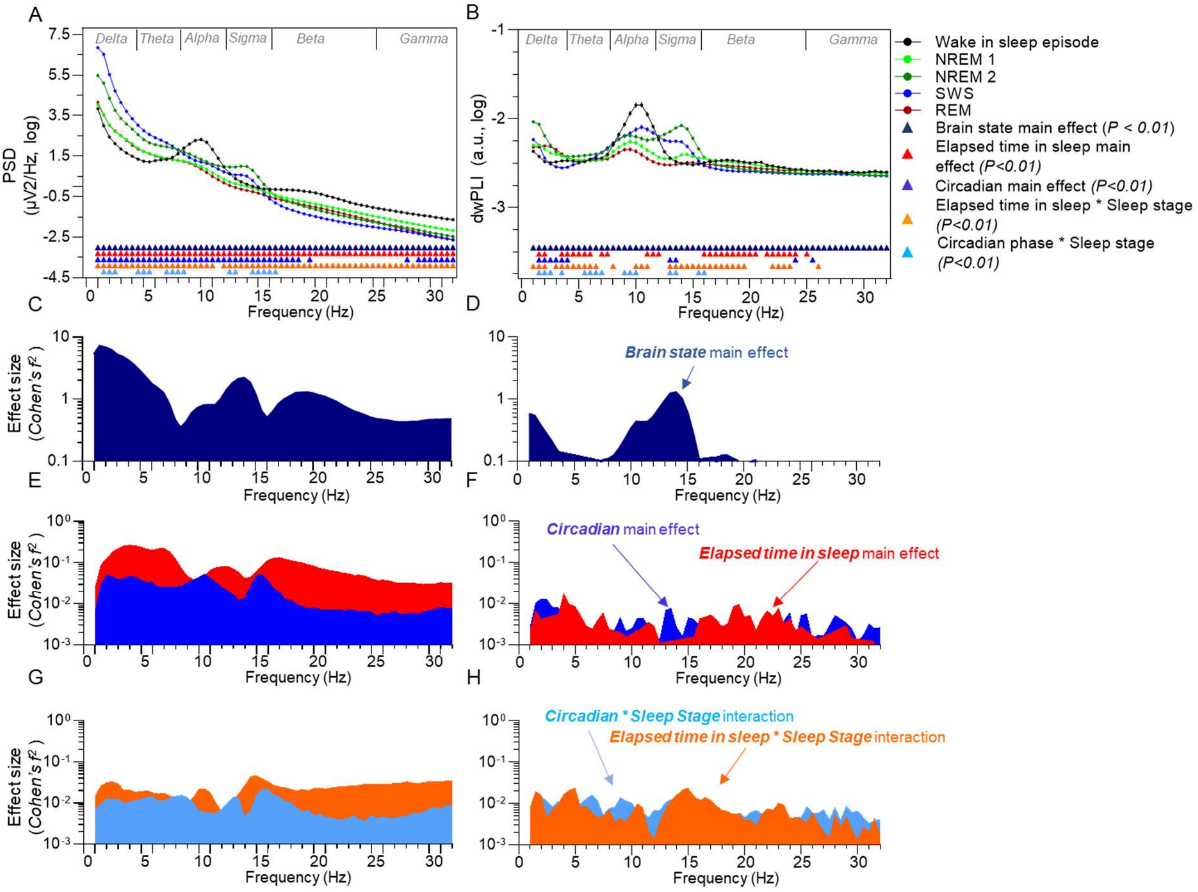


**Fig. S3. Power spectrum density and functional connectivity spectra across brain states and all sleep episodes.** The main effect of vigilance states on (A) power spectrum density (PSD) and (B) debiased weighted Phase Lag Index ($_{dw}$PLI) across all sleep episodes (231 nights) and averaged across EEG channels and cannels pairs, respectively. The upper panels represent least-square means (Lsmeans) and standard error of the mean (SEM). (C and D) Effect size (*Cohen's f*$^2$) of main effects of brain state on the PSD and dwPLI. (E and F) Effect size of main effects of Elapsed time in sleep (red) and Circadian phase (blue) on PSD and dwPLI. (G and H) Effect size of interactions between factors Elapsed time in sleep and Circadian phase on PSD and dwPLI. Colored triangles in (A) and (B) indicate significant (P<.01) effects. Detailed statistical results are in Dataset S2.xlsx**.**

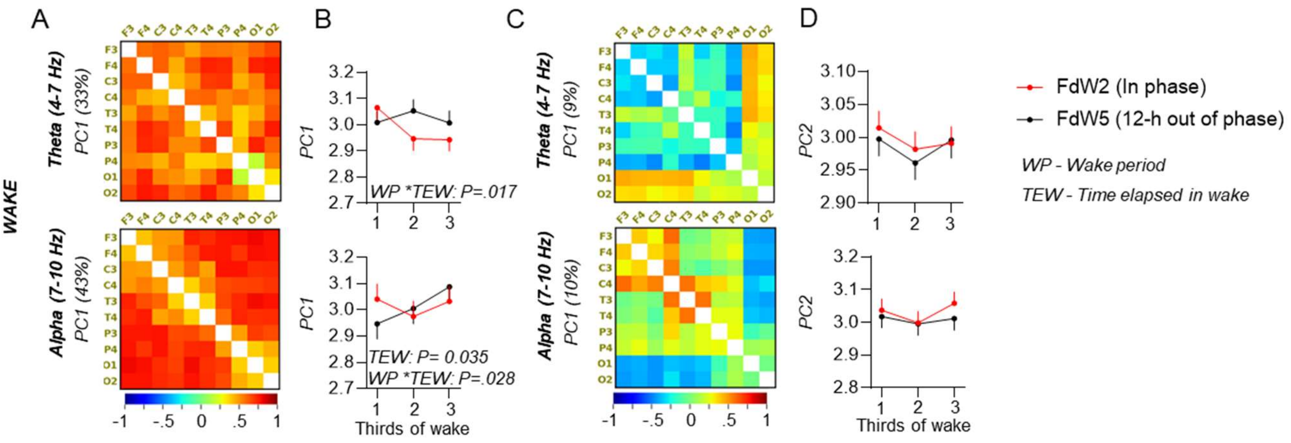


**Fig. S4. Principal component analysis (PCA) of functional connectivity, as measured by the debiased weighted phase lag index (dwPLI), in Wake: time-elapsed-in-wake-dependent and circadian modulation (scheduled wake episode).** Resting EEG was collected during the scheduled wake periods, both before and after each cognitive test session during the Karolinska Drowsiness Test. In total, six cognitive tests were conducted per wake period. (A and C) Loading values of individual electrode pairs on PC1 and PC2, respectively. (B and C) The modulation of PC1 and PC2 by elapsed time spent in wake across the baseline (FdW2) and 12-h-out of phase wake episodes (FdW5) as explained in Fig 1A. Least square means (LSMeans) and standard errors of the mean (SEM) are presented for each 370-minute interval (i.e., one-third of the wake period), across FdW2 and FDW5. Type III fixed effects are reported. Detailed statistical results are in Table S2.

**Supplementary Tables**

| BrainState | FrequencyBand | PC1 | PC2 | PC3 | PC4 | PC5 | PC6 | PC7 | PC8 | PC9 | PC10 |
|---|---|---|---|---|---|---|---|---|---|---|---|
| NREM | Delta | 0.64 | 0.04 | 0.03 | 0.02 | 0.02 | 0.02 | 0.02 | 0.01 | 0.01 | 0.01 |
| | Theta | 0.33 | 0.19 | 0.08 | 0.04 | 0.03 | 0.03 | 0.03 | 0.02 | 0.02 | 0.02 |
| | Alpha | 0.37 | 0.14 | 0.07 | 0.06 | 0.04 | 0.03 | 0.03 | 0.03 | 0.02 | 0.02 |
| | Sigma | 0.38 | 0.10 | 0.06 | 0.05 | 0.04 | 0.04 | 0.03 | 0.03 | 0.02 | 0.02 |
| | Beta | 0.47 | 0.09 | 0.05 | 0.04 | 0.03 | 0.02 | 0.02 | 0.02 | 0.02 | 0.02 |
| | Gamma | 0.49 | 0.05 | 0.04 | 0.03 | 0.03 | 0.03 | 0.02 | 0.02 | 0.02 | 0.02 |
| REM | Delta | 0.44 | 0.08 | 0.05 | 0.05 | 0.04 | 0.03 | 0.03 | 0.02 | 0.02 | 0.02 |
| | Theta | 0.39 | 0.08 | 0.06 | 0.04 | 0.04 | 0.04 | 0.03 | 0.02 | 0.02 | 0.02 |
| | Alpha | 0.43 | 0.09 | 0.06 | 0.05 | 0.04 | 0.04 | 0.03 | 0.02 | 0.02 | 0.02 |
| | Sigma | 0.34 | 0.08 | 0.06 | 0.05 | 0.04 | 0.04 | 0.03 | 0.03 | 0.02 | 0.02 |
| | Beta | 0.38 | 0.07 | 0.05 | 0.05 | 0.04 | 0.03 | 0.03 | 0.03 | 0.03 | 0.02 |
| | Gamma | 0.47 | 0.08 | 0.05 | 0.04 | 0.03 | 0.02 | 0.02 | 0.02 | 0.02 | 0.02 |
| WAKE in Sleep | Delta | 0.35 | 0.08 | 0.05 | 0.04 | 0.04 | 0.03 | 0.03 | 0.02 | 0.02 | 0.02 |
| | Theta | 0.39 | 0.08 | 0.06 | 0.04 | 0.04 | 0.04 | 0.03 | 0.02 | 0.02 | 0.02 |
| | Alpha | 0.48 | 0.09 | 0.08 | 0.06 | 0.06 | 0.03 | 0.03 | 0.02 | 0.02 | 0.02 |
| | Sigma | 0.30 | 0.11 | 0.08 | 0.07 | 0.04 | 0.04 | 0.03 | 0.03 | 0.02 | 0.02 |
| | Beta | 0.32 | 0.09 | 0.06 | 0.05 | 0.04 | 0.04 | 0.04 | 0.02 | 0.02 | 0.02 |
| | Gamma | 0.32 | 0.07 | 0.06 | 0.04 | 0.04 | 0.03 | 0.03 | 0.02 | 0.02 | 0.02 |
| Wake | Delta | 0.39 | 0.08 | 0.06 | 0.05 | 0.04 | 0.03 | 0.03 | 0.03 | 0.02 | 0.02 |
| | Theta | 0.33 | 0.09 | 0.06 | 0.05 | 0.05 | 0.04 | 0.04 | 0.03 | 0.03 | 0.02 |
| | Alpha | 0.43 | 0.10 | 0.06 | 0.05 | 0.04 | 0.03 | 0.03 | 0.03 | 0.02 | 0.02 |
| | Sigma | 0.26 | 0.09 | 0.07 | 0.05 | 0.05 | 0.04 | 0.04 | 0.03 | 0.03 | 0.03 |
| | Beta | 0.42 | 0.15 | 0.06 | 0.04 | 0.03 | 0.03 | 0.03 | 0.02 | 0.02 | 0.02 |
| | Gamma | 0.51 | 0.17 | 0.06 | 0.04 | 0.02 | 0.02 | 0.02 | 0.02 | 0.01 | 0.01 |

**Table S1.** Eigenvalues of the first ten principal components derived from the debiased weighted phase lag index (dwPLI) for delta, theta, alpha, sigma, beta, and gamma frequency bands across NREM, REM, Wake during sleep episodes, and Wake conditions.

| PC-component | Frequency Band | Effect | NumDF | DenDF | FValue | P value | Cohen's $f^2$ |
|---|---|---|---|---|---|---|---|
| **PC1** | **Alpha** | Elapsed Time Awake | 2 | 40.9 | 3.64 | 0.0351 | 0.18 |
| | | Wake Episode | 1 | 9.23 | 0.02 | 0.8877 | 0.00 |
| | | Wake Episode * Elapsed Time Awake | 2 | 40.5 | 3.92 | 0.0278 | 0.19 |
| | **Theta** | Elapsed Time Awake | 2 | 40.2 | 2.37 | 0.106 | 0.12 |
| | | Wake Episode | 1 | 9.77 | 0.72 | 0.4158 | 0.07 |
| | | Wake Episode * Elapsed Time Awake | 2 | 40.1 | 4.52 | 0.017 | 0.23 |
| **PC2** | **Alpha** | Elapsed Time Awake | 2 | 39.3 | 3.09 | 0.0567 | 0.16 |
| | | Wake Episode | 1 | 9.43 | 0.59 | 0.4617 | 0.06 |
| | | Wake Episode * Elapsed Time Awake | 2 | 39.2 | 0.89 | 0.4187 | 0.05 |
| | **Theta** | Elapsed Time Awake | 2 | 43.7 | 1.12 | 0.3364 | 0.05 |
| | | Wake Episode | 1 | 11.4 | 0.56 | 0.4696 | 0.05 |
| | | Wake Episode * Elapsed Time Awake | 2 | 43.5 | 0.17 | 0.8405 | 0.01 |

**Table S2.** Main effects and interactions of elapsed time in wake (by wake-episode thirds) and wake episode condition (habitual timing, FdW2, versus 12 hours out of phase, FdW5) on PC1 and PC2 derived from dwPLI-based functional connectivity in the alpha and theta frequency bands. Type III fixed effects are shown.

| Feature | β OLS | 95% CI (OLS) | P (OLS) | β Linear | β Ridge | β Lasso |
|---|---|---|---|---|---|---|
| Sleep Episode | 0.114 | [-0.17, 0.39] | 0.422 | 0.011 | 0.012 | 0 |
| NREM-Alpha | 0.446 | [0.09, 0.80] | 0.0142 | 0.333 | 0.272 | 0 |
| NREM-beta | 0.604 | [-0.01, 1.22] | 0.0551 | 0.982 | 0.673 | 0 |
| NREM-Delta | -0.116 | [-0.50, 0.27] | 0.5493 | -0.289 | -0.27 | 0 |
| NREM-Gamma | -0.49 | [-0.99, 0.01] | 0.0542 | -0.872 | -0.609 | 0 |
| **NREM-Sigma** | **0.576** | **[0.24, 0.91]** | **0.0008** | 0.668 | 0.619 | 0.261 |
| **NREM-Theta** | **-0.97** | **[-1.30, -0.64]** | **<.00001** | -0.92 | -0.875 | -0.62 |
| REM-Alpha | -0.354 | [-0.66, -0.05] | 0.0217 | -0.421 | -0.395 | 0 |
| REM-Beta | 0.331 | [-0.21, 0.87] | 0.2297 | 0.253 | 0.313 | 0 |
| REM-delta | -0.488 | [-0.86, -0.11] | 0.0109 | -0.706 | -0.57 | 0 |
| REM-gamma | -0.45 | [-0.92, 0.02] | 0.0593 | -0.226 | -0.277 | 0 |
| REM-sigma | -0.052 | [-0.47, 0.37] | 0.8081 | -0.144 | -0.048 | 0 |
| REM-theta | 0.062 | [-0.23, 0.36] | 0.68 | 0.241 | 0.185 | 0 |
| $R^2$ | 0.29 | | | -0.13 | -0.05 | -0.03 |
| Relative RMSE | 0.92 | | | 1.05 | 1.02 | 1 |

**Table S3.** Regression results from multivariate exploratory analyses assessing associations between sleep-related functional connectivity (FC) and cognitive performance**.** The table presents standardized regression coefficients (β), 95% confidence intervals (CI), and p-values from Ordinary Least Squares (OLS) models, alongside coefficients from machine-learning–based predictive models using linear, ridge, and lasso regression. FC was summarized using the first principal component (PC1) extracted from dwPLI-based connectivity matrices across 66 EEG channel pairs per frequency band. The cognitive outcome variable (cognitive PC1) represents a general alertness and working memory factor derived from 21 behavioral measures. Thirteen predictors were entered simultaneously into each model: FC PC1 scores for six NREM (delta, theta, alpha, sigma, beta, gamma) and six REM (delta, theta, alpha, sigma, beta, gamma) frequency bands, as well as the sleep episode (FdN1–FdN7). Model performance is summarized by $R^2$ and relative root mean squared error (Relative RMSE). Bolded OLS coefficients denote predictors with significant associations ($p < 0.01$).

| Brain State | Frequency band | Sleep episode | Spearman Rho | P value |
|---|---|---|---|---|
| NREM | Theta | FdN1 | -0.39 | 0.0211 |
| | | FdN2 | -0.53 | **0.0015** |
| | | FdN3 | -0.26 | 0.1434 |
| | | FdN4 | -0.27 | 0.1415 |
| | | FdN5 | -0.19 | 0.326 |
| | | FdN6 | -0.37 | 0.0416 |
| | | FdN7 | -0.27 | 0.1486 |
| | Sigma | FdN1 | 0.34 | 0.049 |
| | | FdN2 | 0.08 | 0.6653 |
| | | FdN3 | 0.28 | 0.1162 |
| | | FdN4 | 0.15 | 0.4093 |
| | | FdN5 | 0.15 | 0.4346 |
| | | FdN6 | 0.37 | 0.0399 |
| | | FdN7 | 0.30 | 0.106 |

**Table S4.** Summary of Spearman correlations between the first principal component (PC1) of functional connectivity during NREM in each sleep episode and the average cognitive PC1 reflecting general alertness and working memory performance measured six times during the subsequent wake episode. The selected brain states and frequency bands were based on preceding multivariate analyses, which identified only these two NREM frequency bands as significant predictors of cognitive performance, independent of all other predictors.

| feature | β OLS | 95% CI (OLS) | p (OLS) | β Linear | β Ridge | β Lasso |
|---|---|---|---|---|---|---|
| Wake epsiode | -0.132 | [-0.45, 0.19] | 0.41925 | -0.191 | -0.172 | 0 |
| Test Number | -0.484 | [-0.82, -0.15] | 0.0053 | -0.28 | -0.263 | 0 |
| **WAKE_alpha** | **-0.647** | **[-0.99, -0.30]** | **0.00031** | -0.696 | -0.636 | -0.389 |
| WAKE_theta | -0.239 | [-0.59, 0.12] | 0.18513 | 0.072 | 0.054 | 0 |
| $R^2$ | 0.19 | | | -0.13 | -0.13 | -0.17 |
| Relative RMSE | 0.68 | | | 1.05 | 1.05 | 1.07 |

**Table S5.** Regression results from multivariate exploratory analyses assessing associations between wake functional connectivity (FC) and cognitive performance. The table presents standardized regression coefficients (β), 95% confidence intervals (CI), and p-values from Ordinary Least Squares (OLS) models, alongside coefficients from machine-learning–based predictive models using linear, ridge, and lasso regression. FC was summarized using the first principal component (PC1) extracted from dwPLI-based connectivity matrices across 45 EEG channel pairs per frequency band. The cognitive outcome variable (cognitive PC1) represents a general alertness and working memory factor derived from 21 behavioral measures. Four predictors were entered simultaneously into each model: wake FC PC1 scores for the alpha and theta frequency bands, test number (Cog1–Cog6), and wake period (FdW2 vs. FdW5). FC and cognitive performance were measured during the same cognitive test sessions. Model performance is summarized by $R^2$ and relative root mean squared error (Relative RMSE). Bolded OLS coefficients denote predictors with significant associations ($p < 0.01$).

| BrainState | band | Wake episode | Test number | Spearman Rho | P value |
|---|---|---|---|---|---|
| WAKE | Alpha | FdW2 | Cog1 | -0.17 | 0.6021 |
| | | | Cog2 | -0.74 | **0.0058** |
| | | | Cog3 | -0.64 | 0.0353 |
| | | | Cog4 | -0.76 | **0.0062** |
| | | | Cog5 | -0.21 | 0.5372 |
| | | | Cog6 | -0.54 | 0.0709 |
| | | FdW5 | Cog1 | -0.48 | 0.1334 |
| | | | Cog2 | -0.45 | 0.1377 |
| | | | Cog3 | -0.25 | 0.45 |
| | | | Cog4 | -0.56 | 0.0897 |
| | | | Cog5 | -0.73 | 0.0246 |
| | | | Cog6 | -0.71 | 0.0146 |

**Table S6.** Summary of Spearman correlations between the first principal component (PC1) of wake-dependent functional connectivity (FC) in the alpha frequency band and the corresponding cognitive PC1 reflecting general alertness and working memory performance, measured concurrently during each cognitive test session. The alpha band was selected based on preceding multivariate analyses, which identified it as the only frequency band significantly associated with cognitive performance during wakefulness.

### Supplementary Dataset Legends

**Dataset S1.xlsx** – Summary of the main effects and interactions of brain states, elapsed time in sleep, and circadian phase on power spectral density and functional connectivity (FC) spectra across all sleep episodes (231 nights), averaged across EEG channels and channel pairs. Type III fixed effects are reported. Brain states include NREM, REM, and wake; NREM encompasses N2 and slow-wave sleep. Elapsed time refers to time spent in sleep. FC is measured using debiased weighted phase lag index (dwPLI), imaginary coherence (ImCOH), phase coherence (COH), phase lag index (PLI), and weighted phase lag index (wPLI).

**Dataset S2.xlsx** – Summary of the main effects and interactions of brain states, elapsed time in sleep, and circadian phase on power spectral density and FC spectra across all sleep episodes (231 nights), averaged across EEG channels and channel pairs. Type III fixed effects are reported. Brain states include NREM1, NREM2, slow-wave sleep, REM, and wake. FC is measured using dwPLI, ImCOH, COH, PLI, and wPLI.

**Dataset S3.xlsx** – Loading values of individual electrode pairs on PC1 and PC2 across delta, theta, alpha, sigma, beta, and gamma frequencies for NREM, REM, and wake during sleep episodes.

**Dataset S4.xlsx** – Summary of the main effects of elapsed time in sleep and circadian phase on PC1 and PC2 of dwPLI-based FC across delta, theta, alpha, sigma, beta, and gamma frequency bands, and across NREM, REM, and wake. Type III fixed effects are reported.

**Dataset S5.xlsx** – Summary of the main effects of elapsed time in sleep and circadian phase on dwPLI-based FC for each electrode pair across delta, theta, alpha, sigma, beta, and gamma frequency bands, and across NREM, REM, and wake. Type III fixed effects are reported.

**Supplementary Methods**

**Power Spectral Density and Local Connectivity Measures**

All measures were computed as averages of short-time Fourier transforms of artifact-free EEG data, extracted from 4 seconds long windows with 50% overlap, detrended and Hanning tapered. We quantified FC between all pairs of the 12 EEG channels using these Fourier-transforms. Let $x(t)$ and $y(t)$ denote the EEG time series from two electrodes. For each window $w_i$ we computed the Fourier transforms

$$X_i = \mathrm{FFT}(x(w_i)), \quad Y_i = \mathrm{FFT}(y(w_i)) \ .$$

Depending on the particularities of the measure computed from these Fourier transforms as averages over a set of windows, the window number bias can be none, e.g. power spectral density, mild, e.g. coherence, or very strong, e.g. (debiased) weighted phase lag index. Therefore, all measures were estimated from 60 second epochs (N = 29 windows). The length of the epoch was chosen to assure a sufficiently large number of window pairs for the debiased weighted phase lag index (see below). For each 20-minute interval (or 2 minutes for the resting state wake) these estimates were averaged over epochs covering the concatenated artifact-free segments using a 75% (45 seconds) overlap. Our tests on the impact of the fragmentation of the concatenated segments on the extracted measures found no considerable effect.

The connectivity measures are defined as follows:

Coherence captures the consistency of phase and amplitude coupling between $x(t)$ and $y(t)$. It is computed as:

$$\mathrm{Coh} = \frac{\left|\frac{1}{N}\sum_{i=1}^{N} X_i^* Y_i\right|^2}{\left(\frac{1}{N}\sum_{i=1}^{N}|X_i|^2\right)\left(\frac{1}{N}\sum_{i=1}^{N}|Y_i|^2\right)} \ ,$$

where $X_i^*$ is the complex conjugate of $X_i$. Coherence captures total coupling but is susceptible to volume conduction-induced bias, which can inflate connectivity estimates.

**Imaginary coherence** focuses on the imaginary part of the cross-spectrum to mitigate the influence of volume conduction:

$$\mathrm{ImCoh} = \frac{\left|\mathrm{Im}\left(\frac{1}{N}\sum_{i=1}^{N} X_i^* Y_i\right)\right|}{\sqrt{\left(\frac{1}{N}\sum_{i=1}^{N}|X_i|^2\right)\left(\frac{1}{N}\sum_{i=1}^{N}|Y_i|^2\right)}} \ ,$$

where $\mathrm{Im}(\cdot)$ extracts the imaginary component.

**Phase Lag Index (PLI)** measures the consistency of the sign of the imaginary part of the cross-spectrum:

$$\mathrm{PLI} = \left|\frac{1}{N}\sum_{i=1}^{N} \mathrm{sign}\left(\mathrm{Im}(X_i^* Y_i)\right)\right| \ .$$

PLI values near one indicate consistent phase-leading or lagging, while values near zero indicate no consistent phase relationship.

**Weighted Phase Lag Index (wPLI)** extends the PLI by weighting the contribution of each window by the magnitude of its imaginary part:

$$\mathrm{wPLI} = \frac{\left|\sum_{i=1}^{N} \mathrm{Im}(X_i^* Y_i)\right|}{\sum_{i=1}^{N} |\mathrm{Im}(X_i^* Y_i)|} \ .$$

This approach down-weights noisy or near-zero phase differences, increasing robustness.

**Debiased Weighted Phase Lag Index (dwPLI)** corrects for bias due to limited sample size and is computed as:

$$\mathrm{dwPLI} = \frac{\sum_{i\neq j} \mathrm{Im}(X_i^* Y_i)\,\mathrm{Im}(X_j^* Y_j)}{\sum_{i\neq j} \left|\mathrm{Im}(X_i^* Y_i)\,\mathrm{Im}(X_j^* Y_j)\right|} \ ,$$

where the summation runs over all distinct pairs of windows $(i, j)$ with $i \neq j$. In our case N(N - 1)⁄2 = 406 pairs.
